# Sodium ion regulates liquidity of biomolecular condensates in hyperosmotic stress response

**DOI:** 10.1101/2022.06.10.495571

**Authors:** Kazuhiro Morishita, Kengo Watanabe, Isao Naguro, Hidenori Ichijo

## Abstract

Biomolecular condensates are membraneless structures formed through phase separation. Recent studies have demonstrated that the material properties of biomolecular condensates are crucial for their biological functions and pathogenicity. However, the phase maintenance of biomolecular condensates in cells remains elusive. Here, we show that sodium ion (Na^+^) influx regulates the condensate liquidity under hyperosmotic stress. The fluidity of ASK3 condensates increases at the high intracellular Na^+^ concentration derived from extracellular hyperosmotic solution. Moreover, we identified TRPM4 as a cation channel that allows Na^+^ influx under hyperosmotic stress. TRPM4 inhibition causes the liquid-to-solid phase transition of ASK3 condensates, leading to impairment of the ASK3 osmoresponse. In addition to ASK3 condensates, intracellular Na^+^ widely regulates the condensate liquidity and aggregate formation of biomolecules, including DCP1A, TAZ and polyQ-protein, under hyperosmotic stress. Our findings demonstrate that changes in Na^+^ contribute to the cellular stress response via liquidity maintenance of biomolecular condensates.

## Introduction

Cells have multiple compartments called organelles, which enable diverse biological functions. Although classic organelles are sequestered with lipid bilayers to achieve heterogeneous distributions of biomolecules in the intracellular environment, cells also contain numerous membraneless compartments, such as stress granules (SGs) and nucleoli. Recent studies have revealed that many of these membraneless organelles are phase-separated biomolecular condensates, which concentrate specific collections of biomolecules (Banani et al., 2017; Shin and Brangwynne, 2017). Since Brangwynne et al. first proposed the involvement of liquid– liquid phase separation (LLPS) in biological processes (Brangwynne et al., 2009), diverse functions have been suggested for biomolecular condensates; LLPS facilitates the spatiotemporal regulation of biological reactions and subcellular structures (Alberti et al., 2019), which provides one explanation for how biological processes are precisely organized in crowded and complicated intracellular environments (Su et al., 2021). Biomolecular condensates adopt various states with different material properties (e.g., liquid, gel, and solid) depending on intrinsic and extrinsic factors, as well as their generation process (Banani et al., 2017; Boeynaems et al., 2018; Shin and Brangwynne, 2017). Intrinsic factors include genetic mutations and posttranslational modifications of proteins, which change the biomolecules themselves at the atomic level. Extrinsic factors include fluctuations in the environmental conditions, such as temperature, pH and ion concentration, which change the physicochemical state of the whole system. These factors affect the physical interactions between biomolecules; for example, high salt concentrations dissolve condensates or inhibit condensation in vitro (Shin and Brangwynne, 2017) due to the suppression of protein–protein interactions (PPIs) by electrostatic shielding. Hence, biomolecular interactions determine the fluidity of biomolecular condensates, a typical property of the liquid phase derived from the mobility of biomolecules around and across condensates (Bracha et al., 2019).

Recently, growing evidence has suggested the significance of condensate liquidity in their biological functions and pathogenicity, such as selective autophagy, synaptic active zone formation, skin barrier formation, neurodegenerative disease onset and tumorigenesis (Li et al., 2020; McDonald et al., 2020; Molliex et al., 2015; Patel et al., 2015; Peng et al., 2021; Peskett et al., 2018; Quiroz et al., 2020; Watanabe et al., 2021; Yamasaki et al., 2020). In neurodegenerative diseases, genetic alterations such as amyotrophic lateral sclerosis (ALS)-associated mutations in fused in sarcoma (FUS) or heterogeneous nuclear ribonucleoprotein A1 (hnRNPA1) and Huntington’s disease-associated elongations of polyglutamate (polyQ) in Huntingtin accelerate the liquid-to-solid phase transition of each biomolecular condensate, leading to the formation of cytotoxic fibrils (Molliex et al., 2015; Patel et al., 2015; Peskett et al., 2018). Although these intrinsic factors have been well studied under physiological and pathophysiological conditions (Tsang et al., 2020), how condensate liquidity is regulated by extrinsic factors remains elusive in physiological contexts.

When exposed to physicochemical stresses, such as oxidative stress, heat shock and osmotic stress, cells rapidly respond to the induced changes in the intracellular environment to restore cellular homeostasis. These physicochemical stresses can act as extrinsic factors affecting the phase behavior of intracellular biomolecules. For example in the yeast cytosol, heat shock triggers demixing of the poly(A)-binding protein Pab1 (Riback et al., 2017), and pH reduction induces condensation and gelation of the prion protein Sup35 (Franzmann et al., 2018), both of which contribute to cellular fitness during stress. Hence, physiological cellular stresses are suitable study subjects to investigate how extrinsic factors regulate condensate liquidity and how the liquidity maintenance of biomolecular condensates contributes to the cellular stress response. In particular, recent studies have revealed that osmotic stress rapidly induces the condensation of multiple biomolecules (Boyd-Shiwarski et al., 2022; Cai et al., 2019; Carrettiero et al., 2022; Dorone et al., 2021; Jalihal et al., 2020; Watanabe et al., 2021; Yasuda et al., 2020). Osmotic perturbation across the plasma membrane evokes cell swelling or shrinkage following osmotic water influx or efflux, respectively (Hoffmann et al., 2009; Lang et al., 1998). Cell volume changes accompany various alterations in the intracellular environment, including ionic strength, macromolecular crowding and concentrations of biomolecules, all of which could affect the phase behavior of biomolecules (Ishihara et al., 2021). Although several studies have utilized osmotic stress as a means to investigate biomolecules of interest from the perspective of condensates (Cai et al., 2019; Carrettiero et al., 2022; Jalihal et al., 2020; Yasuda et al., 2020), recent studies have demonstrated that osmoresponsive kinases such as apoptosis signal-regulating kinase 3 (ASK3) and with-no-lysine kinase 1 (WNK1) recognize osmotic stress through their intracellular phase transition to mediate signal transduction for volume recovery (Boyd-Shiwarski et al., 2022; Watanabe et al., 2021).

In this study, using ASK3 as a model protein, we demonstrate how the material properties of biomolecular condensates are regulated in cells under hyperosmotic stress. Intracellular sodium ions (Na^+^) regulate the liquidity of ASK3 condensates, and the impairment of the liquidity leads to the abolition of the ASK3 osmoresponse. Our in silico simulation analyses provide mechanistic insight into how the liquidity of ASK3 condensates contributes to ASK3 inactivation under hyperosmotic stress. Furthermore, we show that the liquidity of other biomolecular condensates and the aggregation of polyQ-proteins are also regulated by intracellular Na^+^ under hyperosmotic stress. Our findings demonstrate that changes in intracellular Na^+^ contribute to the cellular stress response via liquidity maintenance of biomolecular condensates.

## Results

### The fluidity of ASK3 condensates increases in high Na^+^ conditions under hyperosmotic stress

Our previous study showed that hyperosmotic stress rapidly induces ASK3 condensation in the cytosol through LLPS regardless of the type of hyperosmotic stress inducer, the solute added for the adjustment of osmolality (Watanabe et al., 2021). However, careful observations of ASK3 condensates revealed that ASK3 condensates move more dynamically and fuse more frequently with each other under sodium chloride (NaCl)-induced hyperosmotic stress than mannitol-induced stress (Video S1). These observational differences were quantified by the morphological changes of ASK3 condensates over time: the count of ASK3 condensates decreased and their mean size increased more rapidly, and the fused ASK3 condensates showed higher circularity under NaCl-induced hyperosmotic stress than mannitol-induced stress (Figure 1A–1C). These morphological characteristics under NaCl-induced hyperosmotic stress are typical of condensates with high fluidity. Thus, we directly examined the fluidity of ASK3 condensates using a fluorescent recovery after photobleaching (FRAP) assay (Bracha et al., 2019). To evaluate the FRAP of dynamically moving condensates, photobleaching was applied to a certain intracellular area that included multiple ASK3 condensates (Watanabe et al., 2021); hence, the FRAP signal in this analysis reflects the overall movements (mobility within a condensate, mobility across the phase boundary, and mobility in the cytosol) of ASK3 molecules in a cell. The FRAP of ASK3 condensates was faster under NaCl-induced hyperosmotic stress than mannitol-induced stress (Figure 1D–1F), suggesting that ASK3 condensates have higher fluidity under NaCl-induced hyperosmotic stress.

**Figure 1.**
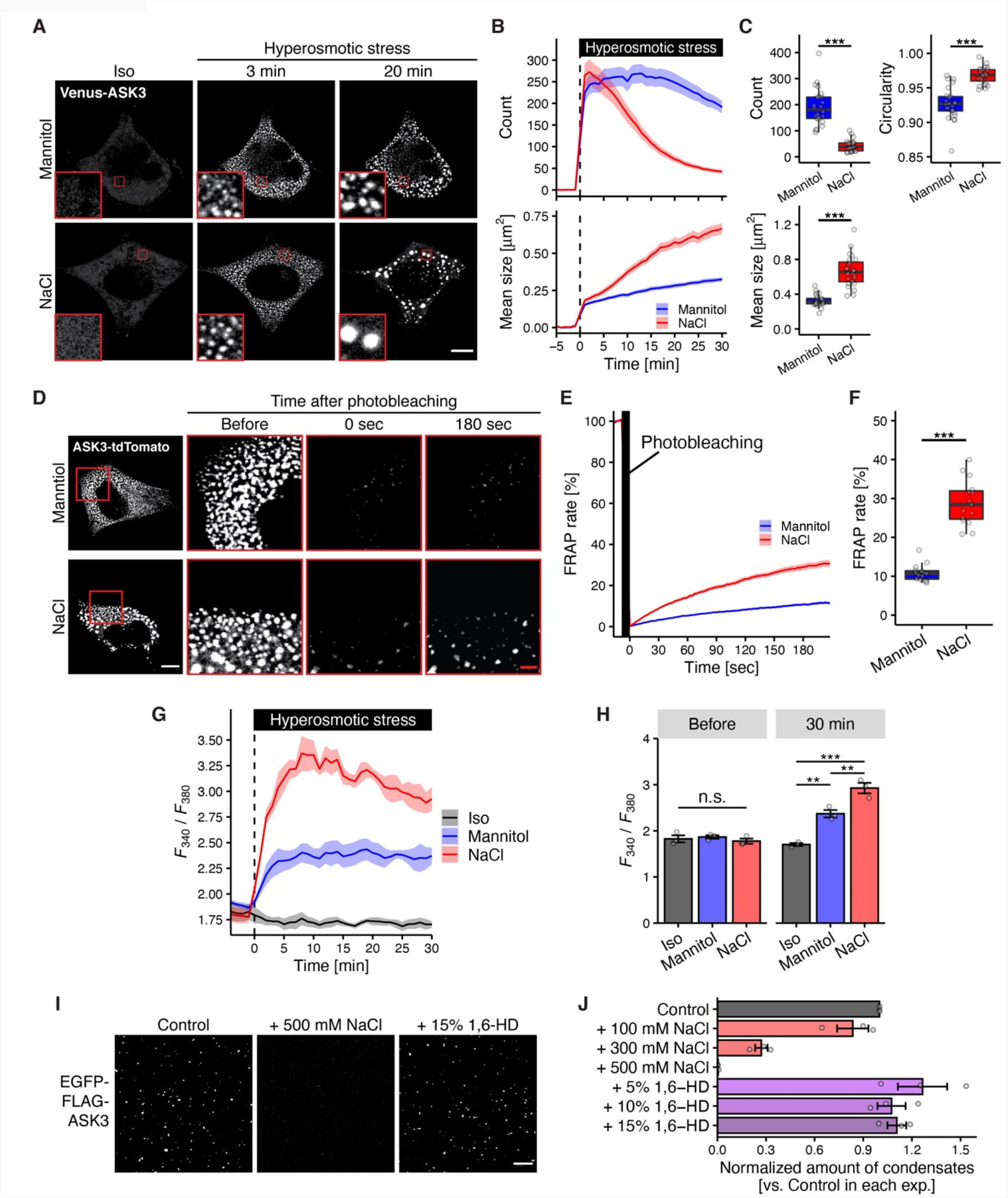
The fluidity of ASK3 condensates increases in high Na^+^ conditions under hyperosmotic stress. (A–C) Morphological changes in ASK3 condensates under hyperosmotic stress. Representative images (A), changes in the count and mean size of ASK3 condensates over time (B), and the count, mean size and circularity at 30 min (C) in Venus-ASK3-HEK293A cells are presented. Hyperosmotic stress: mannitol- or NaCl-supplemented medium (600 mOsm). *N* = 24 cells pooled from 3 independent experiments. ^***^*P* < 0.001 according to two-sided Welch’s *t*-test. Red square: 5 × zoom. (D–F) Fluorescence recovery after photobleaching (FRAP) assay of ASK3 condensates. Representative images (D), changes in the FRAP of ASK3 condensates over time (E) and the FRAP at 180 sec (F) in ASK3-tdTomato-transfected HEK293A cells are presented. Prior to the assay, cells were exposed to hyperosmotic stress (750 mOsm, 20 min). *n* = 15 cells pooled from 3 independent experiments. ^***^*P* < 0.001 according to two-sided Welch’s *t*-test. (G, H) [Na^+^]_i_ under hyperosmotic stress. The change in [Na^+^]_i_ over time (G) and [Na^+^]_i_ at 30 min (H) in HeLa cells are presented. *F*_340_ and *F*_380_ represent fluorescence from SBFI excited at 340 nm and 380 nm, respectively, and *F*_340_/*F*_380_ correlates with [Na^+^]_i_. Hyperosmotic stress: mannitol- or NaCl-supplemented medium (600 mOsm). Data were smoothed by the moving average of 3 time points. *n* = 3 independent experiments. ^**^*P* < 0.01, ^***^*P* < 0.001 according to two-sided unpaired Student’s *t*-tests with Bonferroni correction; n.s. (not significant) according to one-way ANOVA. (I, J) Effects of NaCl and 1,6-hexanediol (1,6-HD) on solid-like ASK3 condensation in vitro. Representative image of fluorescence from EGFP (I) and the quantified amount of solid-like ASK3 condensates (J) are presented. Control: EGFP-FLAG-ASK3 diluted in Tris-buffered saline (TBS) with 1 mM dithiothreitol (DTT) and 10% polyethylene glycol (PEG), 15 min incubation on ice. *n* = 3 independent experiments. Data: mean ± SEM except for boxplots (centerline = median; box limits = [*Q*_1_, *Q*_3_]; whiskers = [max(minimum value, *Q*_1_ − 1.5 × IQR), min(maximum value, *Q*_3_ + 1.5 × IQR)], where *Q*_1_, *Q*_3_ and IQR are the first quartile, the third quartile and the interquartile range, respectively). Scale bar: 10 µm (white) or 2.5 µm (red).

We next investigated what differentiates the fluidity of ASK3 condensates between mannitol- and NaCl-induced hyperosmotic stresses. Given that hyperosmotic stress rapidly induces cation influx from extracellular fluid (Wehner et al., 2003), we hypothesized that the amount of Na^+^ influx could differ between types of hyperosmotic medium even when the final osmolality of the medium is the same. We measured changes in intracellular Na^+^ concentration ([Na^+^]_i_) under hyperosmotic stress using a ratiometric Na^+^ indicator, sodium-binding benzofuran isophthalate (SBFI) (Diarra et al., 2001), which enabled real-time measurements of [Na^+^]_i_ while adjusting the effect of cell volume changes. [Na^+^]_i_ elevated within one minute following hyperosmotic stress, and the elevation was more drastic under NaCl-induced hyperosmotic stress than under mannitol-induced stress (Figure 1G and 1H). Thus, we investigated the possibility that the increased intracellular Na^+^ directly affects the fluidity of ASK3 condensates: Na^+^ itself could affect the fluidity of ASK3 condensates by disrupting excessive PPI with electrostatic shielding. We used a previously developed in vitro assay system for solid-like ASK3 condensates (Watanabe et al., 2021). In this system, NaCl addition inhibited the formation of the solid-like ASK3 condensates, whereas 1,6-hexanediol, a compound that disrupts hydrophobic interactions, did not (Figure 1I and 1J), suggesting that electrostatic interactions perform critical roles in the formation of solid-like ASK3 condensates in vitro. At the same time, because Na^+^ directly contributes to the intracellular osmolality that decides water influx for cell volume recovery (Hoffmann et al., 2009; Lang et al., 1998), we could assume another possibility: the difference in the increased [Na^+^]_i_ between mannitol- and NaCl-induced hyperosmotic stress indirectly resulted in the different fluidity of ASK3 condensates via the difference in cell volume change. In fact, cell volume fluctuation alters intracellular macromolecular crowding, which could also affect the material properties of condensates (Ishihara et al., 2021). We assessed cell volume changes under hyperosmotic stress induced by mannitol or NaCl, using a calcein-quenching system (Hamann et al., 2002). Contrary to the concern, cell volume recovery after NaCl-induced hyperosmotic stress was comparable to or even slower than the recovery after mannitol-induced stress (Figure S1A and S1B), presumably because the overabundance of extracellular Cl^−^ in NaCl-induced hyperosmotic medium prevented Cl^−^ efflux which is necessary for effective cell volume recovery under hyperosmotic stress (Serra et al., 2021). Collectively, these results suggested that intracellular Na^+^ promotes the fluidity of ASK3 condensates in a cell under hyperosmotic stress through electrostatic shielding of excessive ASK3–ASK3 interactions.

### TRPM4 channel activity is necessary for ASK3 inactivation under hyperosmotic stress

For seeking the molecular identity that enables Na^+^ influx to maintain the fluidity of ASK3 condensates, a single cell-level assay is neither robust nor efficient. Given that the liquidity of ASK3 condensates is necessary for ASK3 inactivation under hyperosmotic stress (Watanabe et al., 2021), Na^+^ influx is correspondingly assumed to be necessary for ASK3 inactivation. To investigate this possibility, we evaluated the kinase activity of ASK3 using a specific antibody against phosphorylated Thr808, which is essential for the ASK3 activation (Naguro et al., 2012; Tobiume et al., 2002). ASK3 was inactivated in cells exposed to the hyperosmotic buffer containing Na^+^, while ASK3 inactivation was suppressed in cells exposed to hyperosmotic buffer in which all Na^+^ was replaced with the channel-impermeable cation, *N*-methyl-D-glucamine (NMDG^+^) (Figure 2A). As a route of Na^+^ influx sensitive to hyperosmotic stress, we focused on the hypertonicity-induced cation channel (HICC), which is a monovalent-selective cation channel activated under hyperosmotic stress to facilitate volume recovery (Wehner et al., 2003, 2006). Although its molecular identity remains controversial (Jentsch, 2016), the pharmacological and electrophysiological properties of HICC have been well characterized. Flufenamate is an HICC inhibitor that suppresses Na^+^ influx under hyperosmotic stress (Wehner et al., 2003). We found that flufenamate suppressed ASK3 inactivation under hyperosmotic stress (Figure 2B), suggesting that Na^+^ influx is indeed necessary for ASK3 inactivation. To identify Na^+^ channels activated under hyperosmotic stress, we further characterized the pharmacological sensitivity of endogenous ASK3 inactivation to flufenamate. Endogenous ASK3 activity under hyperosmotic stress was elevated at 5 µM flufenamate (Figure 2C), implying that HICC in HEK293A cells is sensitive to at least 5 µM flufenamate.

**Figure 2.**
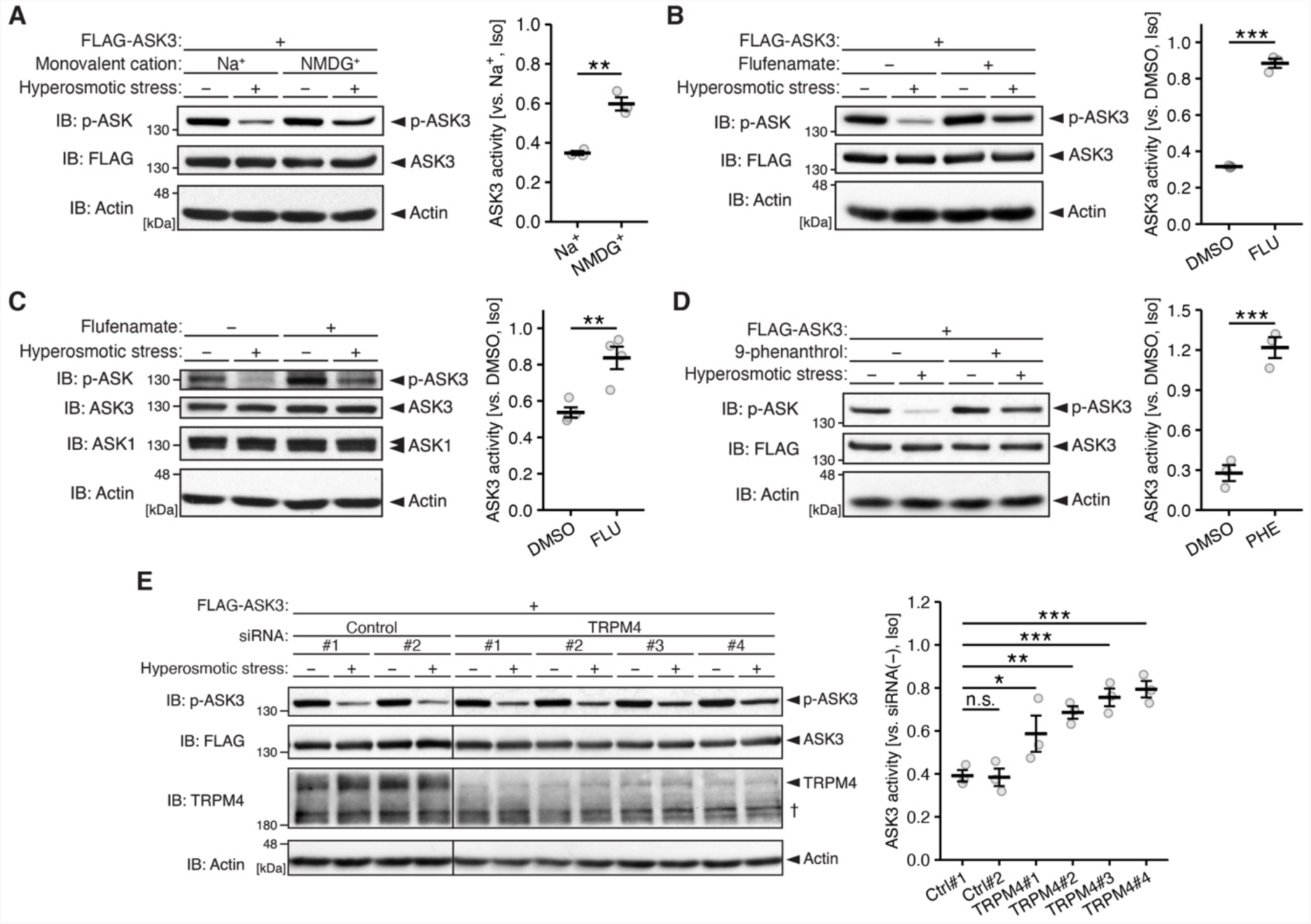
TRPM4 channel activity is necessary for ASK3 inactivation under hyperosmotic stress. (A) Effect of extracellular Na^+^ depletion on ASK3 activity under hyperosmotic stress in FLAG-ASK3-HEK293A cells. Na^+^: osmotic buffer containing 140 mM Na^+^ derived from 130 mM NaCl and 10 mM HEPES-NaOH; NMDG^+^: osmotic buffer in which Na^+^ was replaced with *N*-methyl D-glucamine (NMDG^+^), a channel impermeable cation. (B, C) Effect of HICC inhibition on ASK3 activity under hyperosmotic stress. Flufenamate (−): dimethyl sulfoxide (DMSO), solvent for flufenamate; flufenamate (+): 150 µM (B) or 5 µM (C) flufenamate treated at the same time as osmotic stress. (B) ASK3 activity in FLAG-ASK3-HEK293A cells. (C) Endogenous ASK3 activity in HEK293A cells. (D) Effect of TRPM4 inhibition on ASK3 activity under hyperosmotic stress in FLAG-ASK3-HEK293A cells. 9-phenanthrol (−): DMSO, solvent for 9-phenanthrol; 9-phenanthrol (+): 20 µM 9-phenanthrol treated at the same time as osmotic stress. (E) Effect of TRPM4 depletion on ASK3 activity in FLAG-ASK3-HEK293A cells under hyperosmotic stress. ^†^Nonspecific bands of anti-TRPM4 antibody. TRPM4 was detected in oligomer forms. Quantifications of relative ASK3 activity under hyperosmotic stress are shown to the right of each representative immunoblot. Data: mean ± SEM; *n* = 3 independent experiments. n.s. (not significant), ^**^*P* < 0.01, ^***^*P* < 0.001 according to two-sided unpaired Student’s *t*-tests (A–D) or Dunnett’s test (E). Hyperosmolality (−): 300 mOsm; (+): 400 mOsm; osmolality was adjusted by mannitol; 10 min. IB: immunoblotting.

Although a wide variety of channels have been reported to be inhibited or activated by flufenamate, transient receptor potential melastatin 4 (TRPM4) is the only monovalent-selective cation channel whose IC_50_ to flufenamate is less than 5 µM (Guinamard et al., 2013). TRPM4 is a calcium (Ca^2+^)-activated nonselective monovalent cation channel (Launay et al., 2002) and belongs to the TRP channel superfamily which is proposed to serve as cellular sensors of physicochemical stresses (Venkatachalam and Montell, 2007). Thus far, TRPM4 has been proposed to be involved in osmotic ion/water regulation in astrocytes (Stokum et al., 2018). Furthermore, it was reported that the expression of a dominant negative mutant of TRPM4 suppressed hypertonicity-induced cation influx (Gomi et al., 2014; Launay et al., 2004) and that TRPM4 knockdown showed the tendency to suppress volume recovery following hyperosmotic stress (Koos et al., 2018). Therefore, we assessed TRPM4 as an HICC candidate. 9-Phenanthrol is known as a TRPM4 specific inhibitor (Grand et al., 2008), and we verified that TRPM4 inhibition by the 9-phenanthrol treatment suppressed the elevation of [Na^+^]_i_ (Figure S2A and S2B) and cell volume recovery (Figure S1C and S1D) under hyperosmotic stress. ASK3 inactivation under hyperosmotic stress was inhibited by 9-phenanthrol treatment (Figure 2D), suggesting that Na^+^ influx through TRPM4 is necessary for ASK3 inactivation. To exclude a potential off-target effect of 9-phenanthrol treatment, we also confirmed that TRPM4 depletion using siRNA also suppressed ASK3 inactivation under hyperosmotic stress (Figure 2E). These results suggested that TRPM4 serves Na^+^ influx under hyperosmotic stress, and that Na^+^ influx through TRPM4 contributes to ASK3 inactivation under hyperosmotic stress.

### TRPM4 channel activity maintains the liquidity of ASK3 condensates under hyperosmotic stress

We next investigated whether the fluidity of ASK3 condensates is regulated by TRPM4-dependent Na^+^ influx under hyperosmotic stress. Hereafter, we used 9-phenanthrol for the acute inhibition of TRPM4, enabling suppression of rapid Na^+^ influx following hyperosmotic stress (Figure S2A and S2B). ASK3 condensates showed lower fusion rates, resulting in slower decrease in count and slower increase in mean size under TRPM4 inhibition (Figure 3A–3C). Moreover, the fused ASK3 condensates exhibited lower circularity under TRPM4 inhibition (Figure 3C). From these morphological characteristics, we subsequently examined the fluidity of ASK3 condensates under TRPM4 inhibition and found that ASK3 condensates showed a slower FRAP rate under TRPM4 inhibition (Figure 3D and 3E), suggesting the suppression of ASK3 mobility. We further characterized the material properties of ASK3 condensates under TRPM4 inhibition. As one of their liquid characteristics, ASK3 condensates exhibit reversibility for hyperosmotic stress (Watanabe et al., 2021): ASK3 condensates rapidly disappear after the removal of hyperosmotic stress (DMSO in Figure 3F–3H). However, under TRPM4 inhibition, ASK3 condensates remained after the removal of hyperosmotic stress (Figure 3F–H). Furthermore, the ASK3 condensates remaining under TRPM4 inhibition showed an angular shape with lower circularity, which is the characteristic of solid-like condensates (Figure 3H). Note that the volume recovery under hyperosmotic stress was also suppressed under TRPM4 inhibition (Figure S1C and S1D), which could raise the possibility that the suppressed liquidity of ASK3 condensates under TRPM4 inhibition stemmed from the effects of TRPM4 inhibition on cell volume changes. However, in the analysis of the reversibility of ASK3 condensates, cells completely recovered their volume after the removal of hyperosmotic stress with or without TRPM4 inhibition (Figure S1E and S1F). Collectively, these results suggested that ASK3 condensates undergo liquid-to-solid phase transition in the condition at low [Na^+^]_i_.

**Figure 3.**
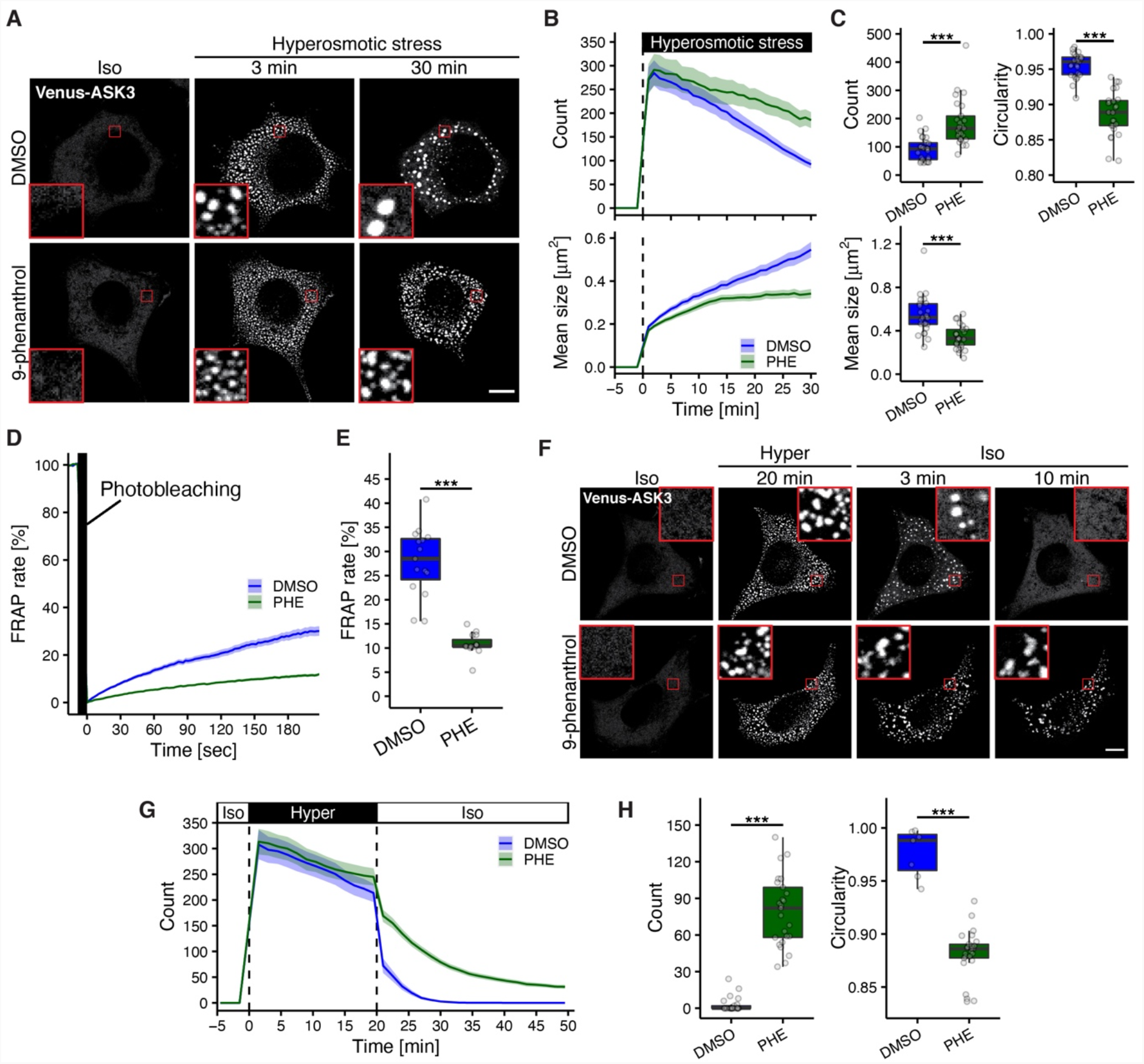
TRPM4 channel activity maintains the liquidity of ASK3 condensates under hyperosmotic stress. (A–C) Morphological changes in ASK3 condensates under TRPM4 inhibition. Representative images (A), changes in the count and mean size of ASK3 condensates over time (B), and the count, mean size and circularity at 30 min (C) in Venus-ASK3-HEK293A cells are presented. *n* = 25 cells pooled from 3 independent experiments. PHE: 20 µM 9-phenanthrol treated at the same time as osmotic stress. Red square: 5 × zoom. (D, E) FRAP assay of ASK3 condensates under TRPM4 inhibition. Changes in the FRAP of ASK3 condensates over time (D) and the FRAP at 180 sec (E) in ASK3-tdTomato-transfected HEK293A cells are presented. Prior to the assay, cells were exposed to hyperosmotic stress (20 min). *n* = 15 cells pooled from 3 independent experiments. PHE: 15 µM 9-phenanthrol treated at the same time as osmotic stress. (F–H) Reversibility of ASK3 condensates under TRPM4 inhibition. Representative images (F), changes in the count of ASK3 condensates over time (G) and the count and circularity at 10 min after the removal of hyperosmotic stress (H) in Venus-ASK3-HEK293A cells are presented. *n* = 25 cells pooled from 3 independent experiments, except for DMSO of the circularity in (H) due to the exclusion of cells with no condensates (*n* = 7 cells pooled from 3 independent experiments). After exposure to hyperosmotic stress for 20 min, osmotic stress was removed by the addition of water to the hyperosmotic medium. PHE: 15 µM 9-phenanthrol treated at the same time as osmotic stress. Red square: 5 × zoom. Hyperosmotic stress: mannitol-supplemented medium (600 mOsm). Data: mean ± SEM except for boxplots (centerline = median; box limits = [*Q*_1_, *Q*_3_]; whiskers = [max(minimum value, *Q*_1_ − 1.5 × IQR), min(maximum value, *Q*_3_ + 1.5 × IQR)], where *Q*_1_, *Q*_3_ and IQR are the first quartile, the third quartile and the interquartile range, respectively). Scale bar: 10 µm. ^***^*P* < 0.001 according to two-sided Welch’s *t*-test.

We previously revealed that ankyrin repeat domain-containing protein 52 (ANKRD52), a subunit of protein phosphatase 6 (PP6) which is an ASK3 phosphatase (Watanabe et al., 2018), forms condensates sharing its phase boundary with ASK3 condensates under hyperosmotic stress (Watanabe et al., 2021) (DMSO in Figure S3). We previously discussed that the partly shared spaces are likely achieved by the liquidity of ASK3 condensates (Watanabe et al., 2021). Thus, we investigated whether TRPM4 inhibition disrupts the partially shared spaces between ASK3 condensates and ANKRD52 condensates. ASK3 condensates shared phase boundaries with ANKRD52 condensates even under TRPM4 inhibition at early time points (∼10 min) after hyperosmotic stress, whereas the partly shared spaces between these two condensates were lost by merging into a single-phase condensate at late time points (Figure S3), further suggesting that Na^+^ influx maintains the liquidity of ASK3 condensates.

### Strength of ASK3–ASK3 interaction regulates condensate liquidity

To obtain insight into the regulatory mechanism for the liquidity of ASK3 condensates by electrostatic shielding, we examined whether the strength of PPI affects the liquidity-derived morphological characteristics of condensates, using the simple computational model for protein diffusion and clustering in a two-dimensional grid space (Dine et al., 2018; Watanabe et al., 2021) (see “General Cluster Model” in the Materials and Methods). Briefly, one protein molecule is regarded as one monomer unit randomly moving around the grid space representing the whole intracellular space of a single cell. To simulate the aggregation-prone behavior of protein units, the model contains a penalty on the movements that accompany detachment from other protein units. This penalty successfully induces cluster formation, which represents biomolecular condensation in cells. In addition to aggregation-prone protein units, obstacle units exist in grid space to reflect intracellular crowded environments with macromolecules (Watanabe et al., 2021). In this model, osmotic cell volume perturbation can be applied by grid space expansion/compression (see the Materials and Methods). Based on the morphological characteristics of ASK3 condensates in cells (i.e., ASK3 condensates were small in count and large in mean size under high [Na^+^]_i_ conditions which provide electrostatic shielding), we speculated that the increased strength of PPI leads to the increase in count and decrease in mean size of clusters in this model. We regarded the parameter of activation energy (Δ*E*) for breaking one interaction between protein units as the strength for PPI. Namely, Δ*E* affects the probability of protein unbinding from clusters (*k*_mov2_) and the protein exchange in clusters (*k*_mov3_), both of which are calculated in accordance with the Arrhenius equation (Figure 4A). When the parameter Δ*E* was increased, the count and mean size of clusters monotonically increased and decreased, respectively (Figure 4B and 4C), due to the decrease in the fusion incidence probability of clusters (Video S2). These results suggested that the liquidity-derived morphological characteristics of clusters are suppressed by tight PPI between protein units. Therefore, this in silico simulation implied that electrostatic shielding contributes to the liquidity of ASK3 condensates by regulating the strength of ASK3–ASK3 interaction.

**Figure 4.**
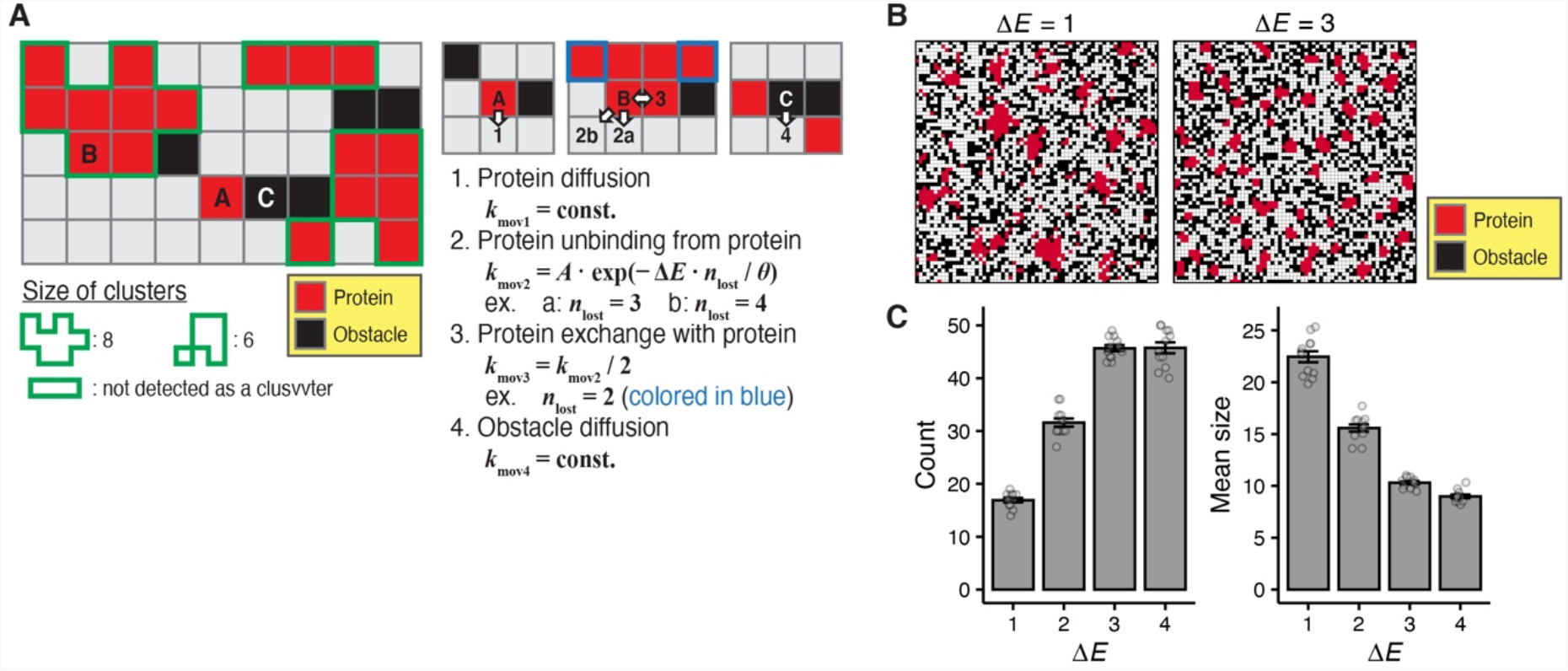
Strength of ASK3–ASK3 interaction regulates condensate fluidity. A computational simulation of the cluster formation of aggregation-prone protein. (A) Schematic diagram of a computational model of protein diffusion and clustering in a two-dimensional grid space. Red square: protein unit, black square: obstacle unit representing intracellular macromolecular crowding, white arrow: potential movement of the selected molecule, Δ*E*: activation energy for breaking one protein–protein interaction (PPI), *n*_lost_: number of interactions broken by the selected movements of the selected protein unit. See the Materials and Methods for the full description of this model. Right three subgrid spaces around molecules A–C are extracted from the left overall diagram. (B, C) A computational simulation for the relationship between the strength of PPI and the count and mean size of clusters. The results after 1 × 10^7^ iteration steps of the rejection kinetic Monte Carlo (rKMC) method for each simulation, representative images of protein clusters under Δ*E* = 1 (weak interaction) or 3 (strong interaction) (B) and the count and mean size of clusters (C) are presented. Data: mean ± SEM, *n* = 12 simulations.

### A decrease in ASK3–ASK3 interaction strength facilitates access to its phosphatase

We further developed the computational model of general aggregation-prone biomolecules into a model specific for ASK3 inactivation by introducing (1) PP6 (ASK3 phosphatase) units, (2) a parameter for the phosphorylation status of ASK3 and (3) ASK3 autophosphorylation and dephosphorylation reaction events (Figure 5A, see “ASK3 Dephosphorylation Model” in the Materials and Methods). Phosphatases also have the aggregation-prone characteristics, which reflects condensation of ANKRD52 in cells under hyperosmotic stress (Figure S3). Based on the observation that ASK3 condensates and ANKRD52 condensates exist in separate phase, regardless of TRPM4 inhibition, at the early time points after hyperosmotic stress when the dephosphorylation of ASK3 was accomplished (Figure 2D and S3), exchange between ASK3 units and PP6 units was not allowed in clusters during simulation. When a PP6 unit is in the destination of phosphorylated ASK3 (p-ASK3) upon movement, or vice versa, p-ASK3 is dephosphorylated at a certain rate. Likewise, dephosphorylated ASK3 (d-ASK3) is autophosphorylated when a d-ASK3 unit is in the destination of p-ASK3, or vice versa. When the grid space was simply shrunk in this model (i.e., the p-ASK3 dephosphorylation rate in clusters was comparable to that outside of clusters), the ratio of p-ASK3 units to total ASK3 (p-ASK3 + d-ASK3) units was not greatly changed (Figure S4A), in contrast to ASK3 inactivation under hyperosmotic stress in cells. For discussion, we previously proposed the hypothesis that ASK3 is efficiently dephosphorylated by PP6 in the phase boundary of ASK3 and PP6 condensates (Watanabe et al., 2021). Therefore, we introduced a constraint that allows the p-ASK3 dephosphorylation event only at ASK3 clusters and observed that the p-ASK3 ratio decreases when the grid space is shrunken in the model with the constraint (Figure 5B), which is consistent with the response of ASK3 activity in cells to hyperosmotic stress. These results suggested the validity of our constrained model as an ASK3 inactivation model and supported our previous hypothesis that p-ASK3 is predominantly dephosphorylated in condensates.

**Figure 5.**
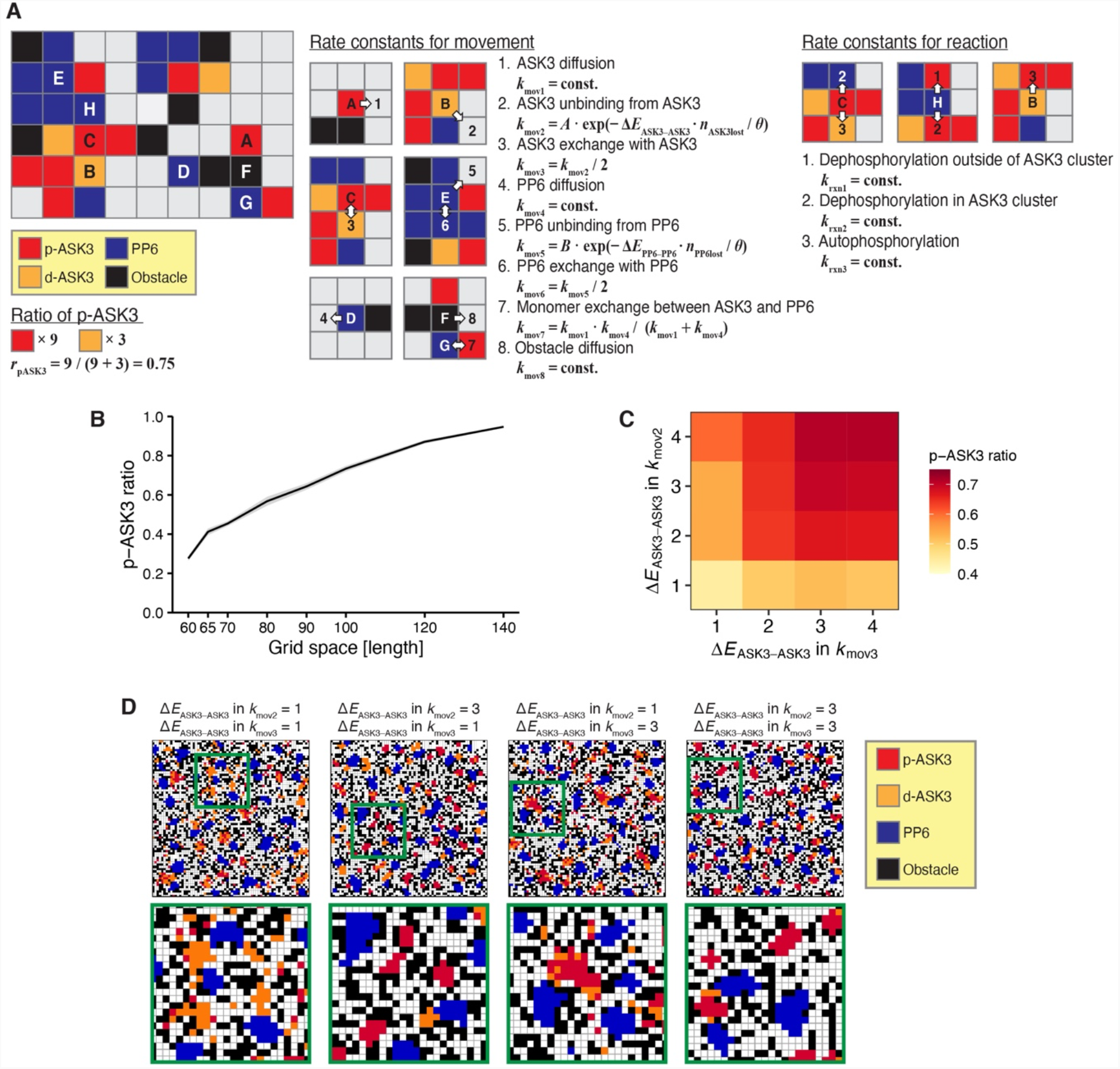
A decrease in ASK3–ASK3 interaction strength facilitates PP6-mediated dephosphorylation of ASK3. (A) Schematic diagram of a computational model for ASK3 inactivation through ASK3 condensation in a two-dimensional grid space. Red square: phosphorylated ASK3 (p-ASK3) unit, orange square: dephosphorylated ASK3 (d-ASK3) unit, blue square: PP6 unit, black square: obstacle unit representing intracellular macromolecular crowding, white arrow: potential movement of the selected molecule. Δ*E*_ASK3–ASK3_: activation energy for breaking one ASK3–ASK3 interaction, *n*_ASK3–ASK3lost_: number of ASK3–ASK3 interactions broken by the selected movements of the selected ASK3 unit. Δ*E*_PP6–PP6_: activation energy for breaking one PP6–PP6 interaction, *n*_PP6–PP6lost_: number of PP6–PP6 interactions broken by the selected movements of the selected PP6 unit, *r*_p-ASK3_: p-ASK3 ratio to total ASK3. The nine subgrid spaces around molecules A–F on the right are extracted from the overall diagram on the left. See the Materials and Methods for the full description of the model. (B) A computational simulation of the relationship between the grid space and p-ASK3 ratio to total ASK3 under the condition that ASK3 is dephosphorylated only at ASK3 cluster. The results after 5 × 10^6^ iteration steps for cluster formation and another 5 × 10^6^ iteration steps for dephosphorylation are presented. Data: mean ± SEM, *n* = 12 simulations. (C, D) A computational simulation of the relationship between the PPI and p-ASK3 ratio. Δ*E*_ASK3–ASK3_ in *k*_mov2_ or *k*_mov3_ was changed independently. A heatmap of the p-ASK3 ratio (C) and representative images for the localization of p-ASK3 and d-ASK3 (D) are presented. (C) Data: mean of *n* = 12 simulations. (D) Lower panels show the 3× zoomed green square regions within upper panels.

Using the established model of ASK3 inactivation, we next investigated how the liquidity of ASK3 condensates contributes to ASK3 inactivation. To clarify how the strength of ASK3– ASK3 interaction affects the phosphorylation status of ASK3, we independently changed Δ*E*_ASK3–ASK3_ in the rate constant for ASK3 unbinding from ASK3 clusters (*k*_mov2_) and that for ASK3 exchange inside clusters (*k*_mov3_), which is one of the advantages of leveraging in silico analysis. When Δ*E*_ASK3–ASK3_ was set to a value corresponding to weak interactions between ASK3 units in both *k*_mov2_ and *k*_mov3_ (i.e., *k*_mov2_ = *k*_mov3_ = 1), ASK3 clusters moved around the grid space and facilitated access to PP6 clusters, and ASK3 units moved dynamically within a cluster, effectively exchanging p-ASK3 with d-ASK3 at the surface region of an ASK3 cluster faced with PP6 clusters (Figure 5C, 5D and Video S3). On the other hand, the increase in Δ*E*_ASK3–ASK3_ of *k*_mov2_ resulted in low morphological flexibility, which hampered ASK3 clusters from easily accessing to the PP6 clusters (Figure 5C and 5D). Furthermore, the increase in Δ*E*_ASK3–ASK3_ of *k*_mov3_ caused the accumulation of dephosphorylated ASK3 at the phase boundary of ASK3 cluster (Figure 5C and 5D), which would interfere with the efficient dephosphorylation of p-ASK3. Additionally, the elevation of Δ*E*_ASK3–ASK3_ in both *k*_mov2_ and *k*_mov3_, which likely corresponds to the actual situation of a cell under TRPM4 inhibition, showed an additive effect on the increase in the p-ASK3 ratio (Figure 5C). In summary, these computational simulations presented one interpretation of how the liquidity of ASK3 condensates contributes to ASK3 inactivation; the liquidity of ASK3 condensates can facilitate ASK3 access to its phosphatase.

### Intracellular Na^+^ regulates condensate liquidity and aggregate formation under hyperosmotic stress

As shown in the computational simulation, the strength of PPI could explain the regulation of condensate liquidity (Figure 4B and 4C). Considering that the General Cluster Model defines only the aggregation-prone characteristics for protein units, we addressed the possibility that not only the liquidity of ASK3 condensates but also that of other biomolecular condensates is generally regulated by [Na^+^]_i_. As model proteins, we selected decapping mRNA 1A (DCP1A), WW domain-containing transcription regulator 1 (WWTR1, also known as TAZ) and WNK1, all of which have been reported to rapidly form condensates following hyperosmotic stress (Boyd-Shiwarski et al., 2022; Cai et al., 2019; Jalihal et al., 2020). DCP1A is commonly known as a marker protein of the processing body (p-body) and forms condensates in the cytosol within seconds under hyperosmotic stress (Figure 6A) (Jalihal et al., 2020). The FRAP of DCP1A condensates was faster under NaCl-induced hyperosmotic stress than under mannitol-induced stress (Figure 6B and 6C). Furthermore, the FRAP of DCP1A condensates was suppressed by TRPM4 inhibition in mannitol-induced hyperosmotic stress (Figure 6D and 6E).

**Figure 6.**
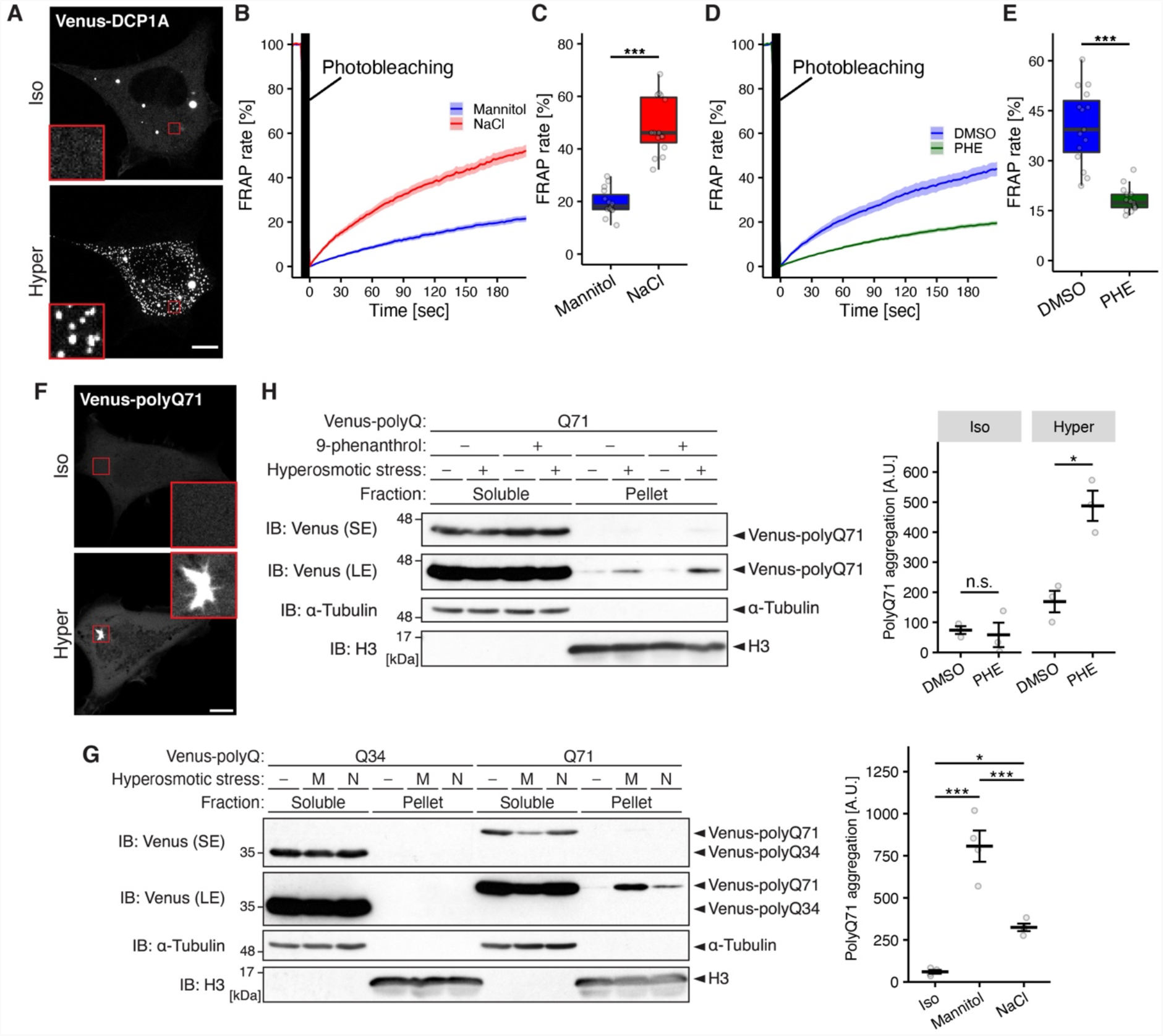
Intracellular Na^+^ regulates condensate liquidity and aggregate formation under hyperosmotic stress. (A) Representative images of DCP1A condensates under hyperosmotic stress. Venus-DCP1A was transfected into HEK293A cells. Hyperosmotic stress: mannitol-supplemented medium (550 mOsm, 30 min). Red square: 5 × zoom. (B–E) FRAP assay of DCP1A condensates. Changes in the FRAP of DCP1A condensates over time (B, D) and the FRAP at 180 sec (C, E) in tdTomato-DCP1A-transfected HEK293A cells are presented. Hyperosmotic stress: mannitol- or NaCl-supplemented medium (600 mOsm) (B, C) or mannitol-supplemented medium (550 mOsm) under TRPM4 inhibition (D, E). PHE: 20 µM 9-phenanthrol treated at the same time as osmotic stress. (F) Representative images of polyQ aggregation under hyperosmotic stress. Venus-polyQ71 was transfected into HEK293A cells. Hyperosmotic stress: mannitol-supplemented medium (600 mOsm, 2 h). Red square: 4 × zoom. (G, H) Triton X-100-insoluble polyQ aggregation under hyperosmotic stress. The Triton X-100 soluble fraction (Soluble) and the Triton X-100 insoluble and SDS soluble fraction (Pellet) were prepared from Venus-polyQ34- or Venus-polyQ71-transfected HEK293A cells. Hyperosmotic stress: mannitol- or NaCl-supplemented medium (600 mOsm, 2 h, represented as M or N, respectively) (G) or mannitol-supplemented medium (500 mOsm, 2 h) under TRPM4 inhibition (H). PHE: 20 µM 9-phenanthrol treated at the same time as osmotic stress. Quantifications of Venus-polyQ71 in the pellet fraction are shown to the right of representative immunoblotting images. α-Tubulin and histone 3 (H3) are shown to ensure the quality of fractionation. SE or LE represents short exposure or long exposure of the electrochemiluminescence, respectively. A. U. represents arbitrary unit. (A, F) Scale bar: 10 µm. (B, D, G, H) Data: mean ± SEM; *n* = 15 cells pooled from 3 independent experiments (B, D), or *n* = 4 (G) or 5 (H) independent experiments. (C, E) Data: centerline = median; box limits = [*Q*_1_, *Q*_3_]; whiskers = [max(minimum value, *Q*_1_ − 1.5 × IQR), min(maximum value, *Q*_3_ + 1.5 × IQR)], where *Q*_1_, *Q*_3_ and IQR are the first quartile, the third quartile and the interquartile range, respectively. n.s. (not significant), ^*^*P* < 0.05, ^***^*P* < 0.001 according to two-sided Welch’s *t*-tests (C, E) or two-sided unpaired Student’s *t*-tests with Bonferroni correction (G, H).

These results suggested that Na^+^ influx regulates the liquidity of DCP1A condensates. Likewise, the Hippo pathway protein TAZ rapidly forms condensates in the cytosol and nucleus under hyperosmotic stress (Figure S5A) (Cai et al., 2019), and TAZ condensates showed the concordant FRAP patterns to ASK3 and DCP1A condensates under these Na^+^ manipulations (Figure S5B–S5E). In contrast to ASK3, DCP1A and TAZ condensates, however, the liquidity of WNK1 condensates was not affected by Na^+^ influx under hyperosmotic stress (Figure S5F–J). In summary, Na^+^ influx regulates the liquidity of not all but multiple hyperosmotic stress-induced biomolecular condensates, which indicates that [Na^+^]_i_ could be a novel environmental factor for regulating the liquidity of intracellular biomolecular condensates.

Based on the observation that ASK3 condensates lost reversibility and underwent liquid-to-solid phase transition in the condition at low [Na^+^]_i_ (Figure 3F–3H), we further explored the possibility that protein aggregation under hyperosmotic stress is also regulated by [Na^+^]_i_. As a model study subject, we focused on polyQ-protein aggregation under hyperosmotic stress (Moronetti Mazzeo et al., 2012). Expansion of the CAG repeat is proposed to augment the formation and the liquid-to-solid transition of polyQ-protein condensates, which likely causes neurodegeneration (Fan et al., 2014; Peskett et al., 2018). While the polyQ-proteins gradually mature into polyQ aggregates over time, hyperosmotic stress quickly induced aggregation-like subcellular structures of a polyQ peptide-conjugated Venus (Venus-Q71), which were observed to display an angular shape (Figure 6F). These polyQ aggregations were detected in the Triton X-100-insoluble pellet fraction under hyperosmotic stress, depending on the polyQ-length (Figure 6G). Venus-Q71 aggregation in the Triton X-insoluble fraction was less induced under NaCl-induced hyperosmotic stress than under mannitol-induced stress (Figure 6G). Furthermore, Venus-Q71 aggregation under mannitol-induced hyperosmotic stress was enhanced by TRPM4 inhibition (Figure 6H). Collectively, these results suggested that intracellular Na^+^ regulates aggregate formation of polyQ-proteins under hyperosmotic stress.

## Discussion

Regulation of the liquidity of biomolecular condensates is critical for their functions and pathogenicity (Li et al., 2020; McDonald et al., 2020; Molliex et al., 2015; Patel et al., 2015; Peng et al., 2021; Peskett et al., 2018; Quiroz et al., 2020; Watanabe et al., 2021; Yamasaki et al., 2020). As the regulatory factors of condensate liquidity, previous studies have presented genetic mutations of proteins and some specific biomolecules, such as ribonucleic acid (RNA), poly(ADP-ribose) and adenosine triphosphate (ATP) (Jain et al., 2016; Roden and Gladfelter, 2021; Tsang et al., 2020; Watanabe et al., 2021). In this study, we demonstrated that Na^+^ influx maintains the liquidity of multiple biomolecular condensates in cells exposed to hyperosmotic stress (Figure 7). From the viewpoint of the cellular osmoresponse, the major role of Na^+^ influx under hyperosmotic stress has been suggested to elevate intracellular osmolality for water influx which eventually leads to cell volume recovery (Hoffmann et al., 2009; Lang et al., 1998). Due to the rapid and drastic changes in the intracellular environment, hyperosmotic stress has the potentially dangerous ability to induce the condensation of many aggregation-prone biomolecules: the elevation of the protein concentration and molecular crowding can cause the phase transition of intracellular environments to an aggregation state in the phase diagram (Figure 7). In fact, the exposure of *Caenorhabditis elegans* to hyperosmotic stress increases the insoluble fraction of extracted proteins (Burkewitz et al., 2011). Additionally, hyperosmotic stress induces the aggregation of neurodegenerative disease-associated proteins (Fragniere et al., 2019; Moronetti Mazzeo et al., 2012), which was suppressed in the condition with higher [Na^+^]_i_ (Figure 6F–6H). Therefore, we speculate that Na^+^ influx can prevent aggregation-prone proteins from falling into an irreversible aggregation state (Figure 7) and thus contribute to proteostasis. Na^+^ is the most abundant charged molecule in extracellular fluids, whereas intracellular Na^+^ is constantly exported to the extracellular side by Na^+^-K^+^-ATPase (Morth et al., 2011). This asymmetrical distribution of Na^+^ and membrane potential achieve a high electrochemical gradient for Na^+^, enabling rapid and substantial Na^+^ influx upon opening of the Na^+^ path. These unique characteristics of Na^+^ physiology present the rationality that cells utilize Na^+^ for the liquidity maintenance of diverse biomolecular condensates under stress responses which require rapid and global counteraction against intracellular perturbation. In summary, Na^+^ influx under hyperosmotic stress would play a pivotal role not only in cell volume recovery but also in maintaining the liquidity of biomolecular condensates.

**Figure 7.**
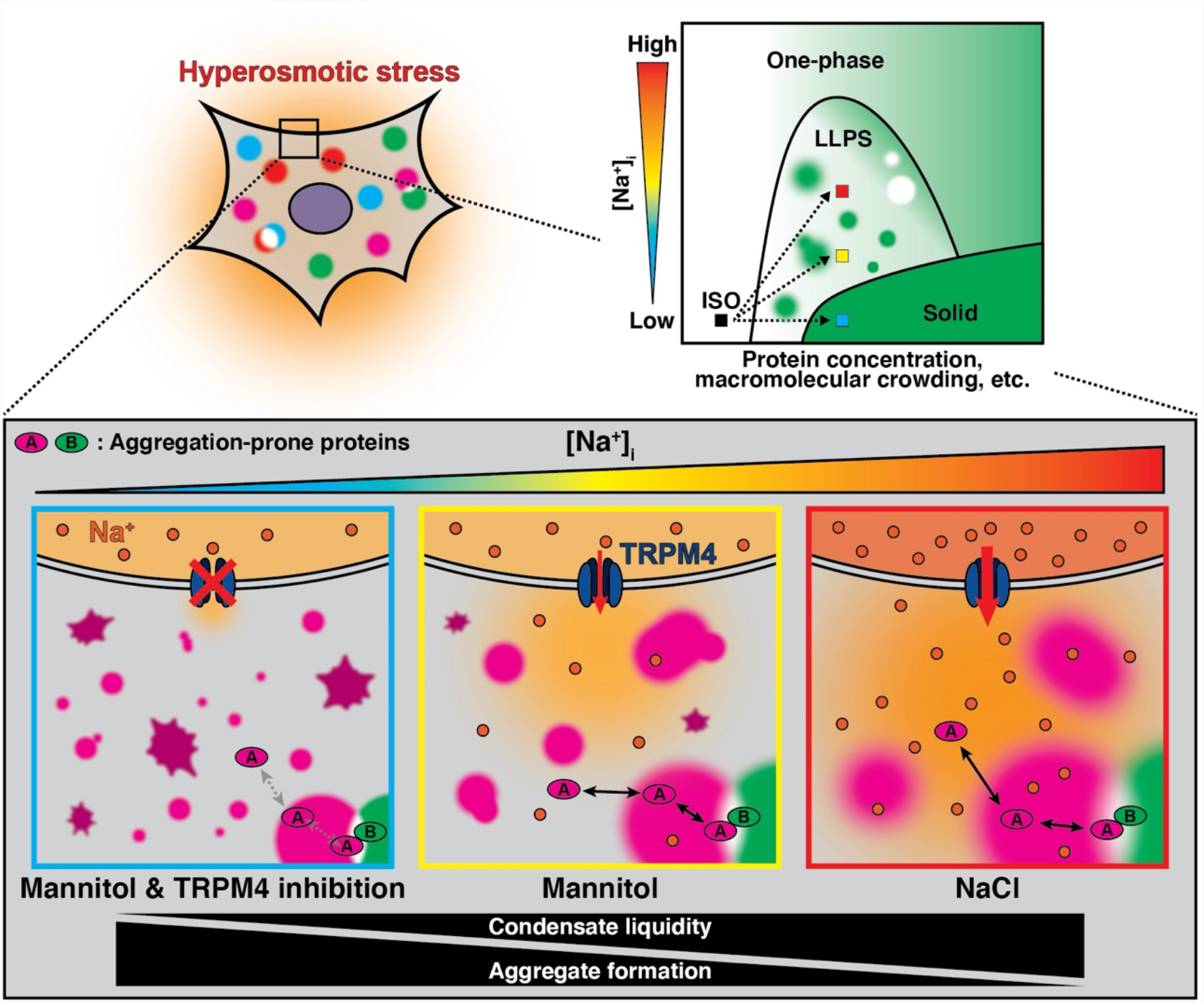
Schematic summary of the main findings in this study. Hyperosmotic stress rapidly changes intracellular environments, such as the elevation of ion/protein concentration and macromolecular crowding. These changes induce the phase transition of biomolecules from a single-phase state to a phase-separated state (dotted arrows in the phase diagram). The elevation of [Na^+^]_i_ by TRPM4 contributes to maintaining condensate liquidity and suppressing aggregate formation. Colored squares in the phase diagram (red, yellow, and cyan) correspond to the color-surrounded conceptual representations in the lower illustration. Under NaCl-induced hyperosmotic stress (red), condensate liquidity is elevated and aggregate formation is suppressed compared to mannitol-induced stress (yellow). Under TRPM4 inhibition (cyan), the condensate undergoes a liquid-to-solid phase transition and aggregate formation is accelerated, which impair biomolecular functions.

Thus far, the effects of salts on condensation and on existing condensates have been well characterized for purified proteins in vitro and for artificial model proteins. NaCl or KCl dissolves condensates (Shin and Brangwynne, 2017), whereas KCl overload induces reentrant of FUS condensation, which might be driven by hydrophobic interactions (Krainer et al., 2021). Additionally, divalent cations, such as Mg^2+^ or Ca^2+^, affect the liquidity and compositions of condensates that are electrostatically mediated by arginine-rich peptides and long single-stranded RNA (Onuchic et al., 2019). Furthermore, divalent heavy metal cations, such as Ni^2+^ and Zn^2+^, trigger condensation and affect the material properties of condensates in artificial client–ligand models (Hong et al., 2020). These effects of ions on the condensates have been suggested to be derived from the electrostatic shielding and/or the specific coordination between protein and cations. However, few studies have addressed the effects of ions on biomolecular condensates in a physiological context due to the difficulties in manipulating intracellular ion concentrations under physiological conditions. Here, we successfully manipulated [Na^+^]_i_ utilizing hyperosmotic stress (Figure 1G, 1H and S2), one of the physiological stresses accompanying intracellular water/ion fluctuations (Hoffmann et al., 2009). [Na^+^]_i_ is estimated to be 5–15 mM under steady-state conditions, whereas the mannitol- or NaCl-supplemented hyperosmotic medium used in this study contained approximately 150 or 300 mM Na^+^, respectively. The difference in the extracellular [Na^+^] under hyperosmotic stress affected Na^+^ influx (Figure 1G and 1H). Additionally, we identified TRPM4 as a cation channel that allows Na^+^ influx under hyperosmotic stress (Figure 2 and S2). Collectively, these manipulations of Na^+^ influx enabled us to investigate the effect of Na^+^ on biomolecular condensates under physiological conditions and to reveal that intracellular Na^+^ regulates the liquidity of multiple condensates (Figure 1, 3 and 6). The sensing mechanism of hyperosmotic stress by TRPM4 remains to be clarified, but it would be worth investigating the involvement of intracellular Ca^2+^ or phosphatidylinositol 4,5-biphosphate (PI(4,5)P_2_) because both rapidly increase under hyperosmotic stress (Dascalu et al., 2000; Yamamoto et al., 2006) and can activate TRPM4 (Launay et al., 2002; Nilius et al., 2006).

We also found a condensate whose liquidity is not affected by [Na^+^]_i_ (Figure S5F–S5J), suggesting that some properties of proteins might be essential for determining the responsiveness to [Na^+^]_i_. Wide varieties of interactions, including electrostatic (dipole–dipole, ionic, cation–π), aromatic (π–π) and hydrophobic interactions, drive condensation and determine material properties of biomolecular condensates (Bracha et al., 2019). Considering that hydrophobic residues were suggested to be crucial for WNK1 condensation (Boyd-Shiwarski et al., 2022), material properties including the liquidity of WNK1 condensates may be regulated primarily by hydrophobic interactions. In contrast, intracellular Na^+^ may regulate the liquidity of biomolecular condensates whose liquid-to-solid transition is regulated mainly by charged amino acids. In fact, WNK1 has a higher ratio of hydrophobic intrinsically disordered regions (IDRs) to charged IDRs (Holehouse et al., 2017; Kyte and Doolittle, 1982; Mészáros et al., 2018) than ASK3, DCP1A or TAZ (Figure S6). Recently, it has been suggested that the amino acid sequence and its composition characteristics of proteins can predict the capacity for condensation using machine learning (Chu et al., 2022; van Mierlo et al., 2021; Saar et al., 2021). When more protein examples regulated by Na^+^ accumulate, a similar machine learning approach will systematically predict whether intracellular Na^+^ regulates the liquidity of condensates composed of uncharacterized proteins, unveiling the molecular grammar of proteins governing phase separation (Wang et al., 2018).

In this study, we showed that the liquid-to-solid phase transition of ASK3 condensates leads to impaired ASK3 osmoresponse. Although recent studies have demonstrated the significance of liquidity maintenance for biomolecular condensates (Li et al., 2020; McDonald et al., 2020; Peng et al., 2021; Watanabe et al., 2021; Yamasaki et al., 2020), few studies have addressed how the mobility of a single biomolecule contributes to its function in condensates. Our in silico analyses provided mechanistic insight into the significance of the condensate liquidity for its own biological functions; both the morphological flexibility of ASK3 condensates and monomer exchangeability within ASK3 condensates, which are derived from the mobility of ASK3 monomers, facilitate ASK3 access to its phosphatase, PP6, at the phase boundary of ASK3 condensates (Figure 5D). Importantly, in silico analyses enabled us to virtually test the two parameters that were difficult to control separately in cellular experiments (Figure 5C and 5D). Recent studies have utilized simple computational simulations to predict the properties of condensates (Dine et al., 2018; Freeman Rosenzweig et al., 2017; Watanabe et al., 2021; Xu et al., 2020). Hence, in silico experiment is a powerful tool to interpret the relationship between single biomolecules and condensate physiology (Alberti et al., 2019).

One of the most important questions derived from our findings is whether intracellular Na^+^ regulates condensate liquidity in physiological contexts other than the hyperosmotic stress response. In fact, in vitro RNA granules showed higher liquidity under high [Na^+^] conditions (Elbaum-Garfinkle et al., 2015; Onuchic et al., 2019); the liquidity of RNA granules may also be regulated by [Na^+^] in intracellular environments. One of the physiological situations that accompany [Na^+^]_i_ dynamics is the neuronal firing. In excitable neurons, the action potential produced by Na^+^ and K^+^ dynamics serves a critical role in neurotransmissions, which accompanies the changes in [Na^+^]_i_ at the local environment around the ion channels. Considering that liquid-to-solid phase transition of condensates at motor neurons and at synaptic active zones are suggested to lead to neuronal dysfunction (McDonald et al., 2020; Patel et al., 2015), the involvement of [Na^+^]_i_ regulation would be worth investigating to elucidate the mechanisms of these toxic aggregations. In summary, further analysis of Na^+^ homeostasis regulation in our body at the point of phase separation would be a clue to understanding aggregation-related diseases.

## Materials and Methods

### Resource availability

#### Lead contact

Further information and requests for resources and reagents should be directed to and will be fulfilled by the lead contact, Isao Naguro.

#### Material availability

Reagents generated in this study will be made available on request.

#### Data and code availability

All data supporting the findings of this study are available within the paper and its Supplementary Information files. Further information and requests for resource and reagents should be directed to I.N.

All custom scripts used in this study are fundamental enough for standard users to code the procedures described in the Materials and Methods, but further information and requests for the custom scripts should be directed to I.N.

### Experimental model and subject details

#### Cell lines and cell culture

HEK293A cells were cultured in Dulbecco’s modified Eagle’s medium (DMEM)-high glucose (Sigma-Aldrich, Cat. #D5796; Wako Pure Chemical Industries, Cat. #044-29765) supplemented with 10% fetal bovine serum (FBS; BioWest, Cat. #S1560-500) and 100 units/mL penicillin G (Meiji Seika, Cat. #6111400D2039). HeLa cells were cultured in DMEM-low glucose (Sigma-Aldrich, Cat. #D6046; Wako Pure Chemical Industries, Cat. #041-29775) supplemented with 10% FBS and 100 units/mL penicillin G. Tetracycline-inducible FLAG-ASK3-stably expressing HEK293A (FLAG-ASK3-HEK293A) cells and tetracycline-inducible Venus-ASK3-stably expressing HEK293A (Venus-ASK3-HEK293A) cells were established previously (Watanabe et al., 2018, 2021). FLAG-ASK3-HEK293A cells and Venus-ASK3-HEK293A cells were cultured in DMEM-high glucose supplemented with 10% FBS, 2.5 µg/mL blasticidin (Invitrogen, Cat. #A1113903) and 50 µg/mL Zeocin (Invitrogen, Cat. #R25001). To induce FLAG-ASK3 or Venus-ASK3, the cells were pretreated with 1 µg/mL tetracyclin (Sigma-Aldrich, Cat. #T7660) 18–24 h before assay. All cells were cultured in 5% CO_2_ at 37°C and verified to be negative for mycoplasma.

### Method details

#### Transfection

Plasmid transfections were performed with polyethyleneimine “MAX” (Polyscience, Cat. #24765) when HEK293A cells were grown to 90% confluency, according to a previously described protocol with minor optimization (Longo et al., 2013). As exceptions, TAZ and WNK1 transfections were performed with Lipofectamine 2000 Transfection Reagent (Invitrogen, Cat. #11668019), according to manufacturer’s instructions. siRNA transfections for FLAG-ASK3-HEK293A cells were carried out by reverse transfection using Lipofectamine RNAiMAX (Invitrogen, Cat. #133778-500) and 10 nM siRNAs (Horizon, ON-TARGET*plus*), according to the manufacturer’s instructions.

#### Osmotic stress treatment

In live-cell imaging, [Na^+^]_i_ measurements and cell volume measurements, osmotic stress was applied by adding 2× osmotic medium into the culture medium, followed by the incubation in 5% CO_2_ at 37°C. For mannitol-induced hyperosmotic stress (∼550, ∼600, ∼700 or ∼750 mOsm/kg H_2_O), DMEM supplemented with 10% FBS and 500, 600, 800 or 900 mM mannitol was used as the 2× hyperosmotic medium. In the case of NaCl-induced hyperosmotic stress (∼600, ∼700 or ∼750 mOsm/kg H_2_O), DMEM supplemented with 10% FBS and 300, 400 or 450 mM NaCl was used as the 2× hyperosmotic medium. For the removal of hyperosmotic stress, ultrapure water was added into the ∼600 mOsm hyperosmotic medium to be isoosmotic condition (∼300 mOsm). For TRPM4 inhibition, 9-phenanthrol (Sigma-Aldrich, Cat. #211281) dissolved in dimethyl sulfoxide (DMSO; Sigma-Aldrich, Cat. #D5879) was added to 2× or 3× hyperosmotic medium by 1:1,000 or 1:2,000 dilution.

In immunoblotting experiments, osmotic stress was applied by exchanging the culture medium with osmotic buffer. The isoosmotic buffer (300 mOsm/kg H_2_O, pH 7.4) contained 130 mM NaCl, 2 mM KCl, 1 mM KH_3_PO_4_, 2 mM CaCl_2_, 2 mM MgCl_2_, 10 mM 4-(2-hydroxyethyl)-1-piperazineethanesulfonic acid, 10 mM glucose and 20 mM mannitol. The hyperosmotic buffer (400 mOsm/kg H_2_O, pH 7.4) was the same as the isoosmotic buffer except containing 120 mM mannitol. Na^+^ was completely replaced with NMDG^+^ (Wako Pure Chemical Industries, Cat. #132-08012) in Na^+^-replaced isoosmotic and hyperosmotic buffer. For HICC or TRPM4 inhibition, flufenamate (*N*-(3-trifluoromethylphenyl)anthranilic acid; Tokyo Chemical Industry, Cat. #T2354) or 9-phenanthrol dissolved in DMSO was added to osmotic buffer, respectively. As an exception, osmotic stress was applied by adding 2× hyperosmotic medium in polyQ aggregation experiments.

Absolute osmolality was verified by an Osmomat (Gonotec) osmometer to fall within the range of 295 to 320 mOsm/kg H_2_O for isoosmotic buffer or ±20 mOsm/kg H_2_O for hyperosmotic medium or hyperosmotic buffer. The difference in the osmolality of mannitol-induced hyperosmotic medium and NaCl-induced one was within 5 mOsm in the identical experiments. The difference in the osmolality of normal hyperosmotic buffer and Na^+^-replaced one was within 5 mOsm in the identical experiments.

#### Live-cell imaging

Cells were seeded in a 35 mmφ glass bottom dish (Matsunami, Cat. #D11130H) which were coated with 1% gelatin (Nacalai Tesque, Cat. #16605-42) in phosphate-buffered saline (PBS; 137 mM NaCl, 3 mM KCl, 8 mM Na_3_PO_4_·12H_2_O, 15 mM KH_2_PO_4_) in advance. For transfected HEK293A cells, the cells were reseeded from a 24-well plate into glass bottom dishes 18–30 h after transfection. After 40–56 h, the culture medium was replaced with 1 mL culture medium per dish, and the dish was subsequently viewed by TCS SP5 (Leica) confocal-laser scanning microscope equipped with a stage top incubator (Tokai Hit). The cells were observed in 5% CO_2_ at 37°C using an HC PL APO 63×/1.40 oil objective (Leica). Single- or multi-channel time-lapse images were acquired in 4–8 fields with 4 averages per frame in 1-, 1.5- or 10-min intervals. Venus or tdTomato was excited at 514 nm with argon laser or at 561 nm with a DPSS laser, respectively, and detected by HyD detector (Leica). After obtaining image sets for 5 min under isoosmotic conditions as the “Before” condition, the cells were exposed to osmotic stress by adding 1 mL of 2× osmotic medium per dish and continuously observed for 30 min. Of note, although the cellular morphology was appreciably changed under osmotic stress, we observed the constant position, XY and focal plane, using a motorized stage and the on-demand mode of adaptive focus control system (Leica) in each field.

In the representative movies of the dynamics and fusion of ASK3 condensates, single-channel time-lapse imaging for Venus was performed in a single field with 2 averages per frame at the minimum intervals (∼1 s). To hold a constant position and minimize the autofocusing time, the continuous mode of adaptive focus control system was applied.

In the experiments of the reversibility of ASK3 condensates, single-channel time-lapse images for Venus were captured by the following procedure. After acquiring “Before” image sets for 1–5 frame under culture medium, the cells were exposed to hyperosmotic stress (600 mOsm) by adding 1 mL of 2× hyperosmotic medium per dish and observed for 20 min. Subsequently, the cells were reverted to isoosmotic conditions by adding 2 mL of ultrapure water per dish and observed for another 30 min.

For presentation, representative raw images were adjusted in brightness and contrast linearly and equally among the samples by using Fiji/ImageJ software (Schindelin et al., 2012). Of note, we applied adjustments in the brightness and contrast to multichannel time-lapse image for the ASK3-tdTomato and ANKRD52-Venus in each sample separately; therefore, the signal intensity cannot be compared between samples. To create a time-lapse video, a series of images were equally adjusted in brightness and contrast, captions were added, and the images were converted to a movie file using Fiji/Image software.

For quantification, we used macro scripts for Fiji/ImageJ shown in the previous study to calculate the count and mean size of ASK3 condensates in a cell per frame, and applied it to all raw image sets in batch mode (Watanabe et al., 2021). Briefly, based on the intensity from Venus, the region of interest (ROI) was first defined as the whole cell area of a main cell because there are condensates from other cells in some cases. After applying Gaussian filter, the Venus signal within the ROI was subsequently extracted from a Venus image in accordance with the local threshold. Finally, particle analysis was performed to quantify the number, mean size and circularity of ASK3 condensates. Circularity in this study was defined as 4*πS*/*L*^2^, where *S* and *L* represent area and perimeter of condensates, respectively. Each parameter was determined from pilot analyses in Venus-ASK3-HEK-293A cells. The exported data table was summarized with RStudio software. The Fiji/ImageJ scripts also exported images of both ROIs and identified particles, enabling us to confirm the quality. We excluded several datapoints from the data analysis: (1) if the image was out-of-focus or (2) if the target cell was shrunken or swollen to detach completely.

#### Fluorescent recovery after photobleaching (FRAP) assay

For the quantification of the FRAP of dynamically moving condensates, we applied the FRAP procedure used in the previous study (Watanabe et al., 2021). Prior to FRAP assay, tdTomato-tagged constract-transfected HEK293A cells were placed under the microscope in 5% CO_2_ at 37°C and exposed to hyperosmotic stress (550, 600, 700 or 750 mOsm) for 15 min, which makes the size of condensates ideal for the assay. Subsequently, single-channel time-lapse imaging for tdTomato was performed in a single field with 4 average per frame with a minimum interval (∼2.1 s) using the continuous mode of adaptive focus control system. After acquiring 5 frames as the “Before” condition, a rectangular area that included more than ten condensates was photobleached by the maximum intensity of the DPSS laser for 3 frames, followed by the time-lapse imaging of 100 intervals as the “After” condition. Note that intensely severe hyperosmotic stress was used in Figure 1D–1F because ASK3 condensates move too dynamically under milder NaCl-induced hyperosmotic stress to be tracked for the quantification. Thus, mannitol-induced hyperosmotic stress was used in the experiments of TRPM4 inhibition (Figure 3D and 3F) to analyze the FRAP rate of ASK3 condensates under milder hyperosmotic condition. Accordingly, mannitol-induced hyperosmotic stress was used in the other experiments accompanying TRPM4 inhibition or inhibition of Na^+^ influx (Figure 2, 3, 6D, 6E, 6I and 6J).

To quantify the FRAP rate of ASK3 condensates from image data, particle tracking analysis was first executed for all condensates in an image by using a Fiji plugin TrackMate (Tinevez et al., 2017). In TrackMate, each condensate was identified in each frame by Laplacian of Gaussian detector, followed by connecting frames by linear assignment problem tracker. Each parameter was determined from pilot analysis for tdTomato-tagged each protein. In this tracking analysis, we excluded the condensates (1) that were not successfully tracked from “Before” to “After” or (2) that were present in less than 25 frames. Next, tracking data table was systemically calculated to the FRAP rate in RStudio software. In the R script, each of tracked condensates was first categorized into two groups, photobleached or not-photobleached, based on the XY coordinates of the photobleached rectangular area. Meanwhile, the fluorescence intensity value of condensate *i* at time *t, F*_*i*_(*t*) was converted the relative fluorescence change *F*_*i*_(*t*)/*F*_*i*,Before_, where *F*_*i*,Before_ indicates the mean of *F*_*i*_(*t*) for each condensate *i* in the “Before” condition. At this step, we established a few false positive condensates in the photobleached group whose *F*_*i*_(*t*)/*F*_*i*,Before_ did not exhibit at least a 15% decrease between the “Before” and “After” conditions, whereas *F*_*i*_(*t*)/*F*_*i*,Before_ of the other photobleached condensates dropped by an average of ∼80% in our FRAP assay. To correct the quenching effects during observation, each *F*_*i*_(*t*)/*F*_*i*,Before_ in the photobleached group was normalized to *G*_*i*_(*t*) = (*F*_*i*_(*t*)/*F*_*i*,Before_)/(the mean of *F*_*i*_(*t*)/*F*_*i*,Before_ in the nonphotobleached group). To mitigate the effect of condensate movement in a direction vertical to the focal place on the changes in fluorescence, *G*_*i*_(*t*) was further converted to the mean of *G*_*i*_(*t*) in the photobleached group, *G*(*t*); namely, we summarized all values of photobleached condensates in a cell into representative values of one virtual condensate. Finally, the FRAP rate [%] at time *t* in the cell was calculated as (*G*(*t*) – *G*_Min_)/(1 – *G*_Min_) × 100, where *G*_Min_ was the minimum value within the first three time points of the “After” condition.

#### In vitro condensation assay

For protein purification, HEK293A cells were seeded in 10 cmφ dishes and transfected with EGFP-FLAG-tagged construct. After washing with PBS, the cells were lysed in lysis buffer (20 mM Tris-HCl pH7.5, 150 mM NaCl, 5 mM ethylene glycol-bis(2-aminoethylether)-*N,N,N’,N’*-tetraacetic acid (EGTA), 1% sodium deoxycholate, 1% Triton X-100 and 12 mM β-glycerophosphatase) supplemented with protease inhibitors (1mM phenylmethylsulfonyl fluoride (PMSF) and 5 µg/mL leupeptin), phosphatase inhibitor cocktail II (8 mM NaF, 12 mM β-glycerophosphatase, 1 mM Na_3_VO_4_, 1.2 mM Na_2_MoO_4_, 5 µM cantharidin and 2 mM imidazole) and 1 mM dithiothreitol (DTT). The cell extracts were collected with a scraper from 3 dishes into a single microtube for each protein, followed by centrifugation at 4°C and ∼17,500 × *g* for 10 min. The supernatants were incubated with anti-FLAG antibody beads (Sigma-Aldrich, clone M2, Cat. #A2220) at 4°C for ∼3 h. The beads were washed 4 times with wash buffer (20 mM Tris-HCl pH 7.5, 500 mM NaCl, 5 mM EGTA, 1% Triton X-100 and 2 mM DTT) and once with TBS (20 mM Tris pH 7.5, 150 mM NaCl and 1 mM DTT). The EGFP-FLAG-tagged proteins were eluted from the beads with 0.1 mg/mL 3x FLAG peptide (Sigma-Aldrich, Cat. #F4799) in TBS at 4°C for more than 1 h, followed by dilution to 40 µM with TBS. The concentration of the protein was estimated from the absorbance at 280 nm measured by a SimpliNano (GE healthcare) microvolume spectrophotometer with the extinction coefficient calculated by using the ExPASy ProtParam tool (https://web.expasy.org/protparam/).

The purified EGFP-FLAG-tagged protein was diluted into a sample in a microtube, whose control conditions were 10 µM EGFP-FLAG-tagged protein, 150 mM NaCl, 20 mM Tris (pH 7.5), 20% PEG 4000 (Kanto Kagaku, Cat. #32828-02) and 1 mM DTT. For the inhibition of electrostatic or hydrophobic interactions, NaCl or 1,6-hexanediol (Wako Pure Chemical Industries, Cat. #081-00435) was included in a sample at the indicated concentration, respectively.

The prepared sample was subsequently incubated at 4°C for 15 min. The reaction mixture was immediately loaded into a counting chamber with a cover slip (Matsunami, Cat. #C018241) followed by observation using a TCS SP5 microscope with a 63×/1.40 oil objective. To maintain a constant focal plane even if there were no condensates, we set the focal plane adjacent to the surface of the cover slip using the motorized stage and the on-demand mode of adaptive focus control system. Images of the EGFP signal were captured from 5 random fields per sample. Of note, we began from the protein purification in each independent experiment.

For presentation, representative raw images were adjusted in brightness and contrast linearly and equally within the samples using Fiji/ImageJ software. For quantification, we used macro scripts for Fiji/ImageJ shown in the previous study to calculate the fluorescence intensity and area of ASK3 condensate in each sample and applied the script to all raw images set in batch mode (Watanabe et al., 2021). In the script, a Gaussian filter and background correction were applied to each image, followed by particle analysis. Each parameter was determined from pilot analyses for EGFP-FLAG-ASK3 WT. The exported data table was further summarized in RStudio software. In the R script, the amount of ASK3 condensates in a sample was defined as the mean of total intensity within 5 fields. The amount of condensates was further converted to the relative to the mean of control sample for each experiment.

#### [Na^+^]_i_ measurements

[Na^+^]_i_ was measured using Na^+^ indicator, sodium-binding benzofuran isophthalate (SBFI). SBFI enables ratiometric measurements of [Na^+^] (Diarra et al., 2001), which is suitable for the [Na^+^]_i_ measurements under osmotic volume perturbation accompanying the changes in the intracellular concentration of the indicator. Fluorescence by the excitation at 380 nm (*F*_380_) decreases with the increase in [Na^+^], whereas fluorescence by the excitation at 340 nm (*F*_340_) is not affected by [Na^+^]; *F*_340_/*F*_380_ positively correlates with [Na^+^]_i_.

HeLa cells were seeded in a 96-well plate (BM Bio, Cat. #215006) and incubated for 48 h. Prior to measurements, the mixture of 20 µM SBFI-AM (Invitrogen, Cat. #S1263) and Pluronic F-127 (Invitrogen, Cat. #P3000MP) was loaded via medium change, and the plate was incubated in 5% CO_2_ at 37°C for 4 h, followed by two medium changes with isoosmotic buffer for washing and then with isoosmotic culture medium at 100 µL per well. Before applying hyperosmotic stress, fluorescence (Ex: 340 ± 9 or 380 ± 9 nm, Em: 510 ± 20 nm) was measured in 5% CO_2_ at 37°C for 5 min at 1-min intervals using an Infinite 200 Pro (Tecan). Then, the cells were exposed to hyperosmotic stress (600 mOsm) by adding 100 µL of 2× hyperosmotic medium per well, and subsequent measurements were performed in 5% CO_2_ at 37°C for 30 min at 1-min intervals.

For the data analysis, each fluorescent value was applied background subtraction by the fluorescent value of non-SBFI-loaded well, followed by the calculation of the ratio at time *t, R*(*t*) = *F*_340_(*t*)/*F*_380_(*t*). To smooth the fluctuations of *R*(*t*), the moving average of 3 time points (*R*_rolling_(*t*) = (*R*(*t*−1) + *R*(*t*) + *R*(*t*+1))/3) was presented in the figures. In the case of TRPM4 inhibition, *F*_380_ alone was used for the estimation of [Na^+^]_i_ because 340 nm wavelength was absorbed by 9-phenanthrol. Following background subtraction, the relative fluorescence change to the mean of fluorescent values before hyperosmotic stress, *F*_380_(*t*)/*F*_380, before_, was calculated. For the drift correction of [Na^+^]_i_-independent quenching of SBFI and the cellular response against osmotic stress-operation, *F*_380_(*t*)/*F*_380, before_ was normalized by *F*_380_(*t*)/*F*_380, before_ of 300 mOsm at each time point: *G*(*t*) = {*F*_380_(*t*)/*F*_380, before_}/{*F*_380_(*t*)/*F*_380, before_ of 300 mOsm}. This *G*(*t*) value negatively correlates with [Na^+^]_i_ at time *t*, thus the transition of [Na^+^]_i_ was estimated by its reciprocal, *H*(*t*) = 1/*G*(*t*). To clarify the transition of *H*(*t*), the moving average of 3 time points (*H*_rolling_(*t*) = (*H*(*t*−1) + *H*(*t*) + *H*(*t*+1))/3) was presented in the figures.

#### Immunoblotting

For experiments of ASK3 inactivation, cells were lysed in lysis buffer (20 mM Tris-HCl pH7.5, 150 mM NaCl, 10 mM EDTA, 1% sodium deoxycholate, and 1% TritonX-100) supplemented with protease inhibitors (1 mM PMSF and 5 µg/mL leupeptin). When detecting the phosphorylation of endogenous protein, phosphatase inhibitor cocktail II (20 mM NaF, 30 mM β-glycerophosphatase, 2.5 mM Na_3_VO_4_, 3 mM Na_2_MoO_4_, 12.5 µM cantharidin and 5 mM imidazole) was also supplemented. Cell extracts were clarified by centrifugation at 4 °C and ∼17,500 × *g* for 10 min, and the supernatant was sampled by adding 2× sample buffer (80 mM Tris-HCl pH 8.8, 80 µg/mL bromophenol blue, 28.8% glycerol, 4% sodium dodecyl sulfate (SDS) and 20 mM DTT), followed by boiling at 98°C for 3 min.

For experiments of polyQ aggregation, cells were lysed in lysis buffer (20 mM Tris-HCl pH7.5, 150 mM NaCl, 10 mM EDTA, and 1% TritonX-100) supplemented with protease inhibitors (1 mM PMSF and 5 µg/mL leupeptin). Cell extracts were clarified by centrifugation at 4 °C and ∼17,500 × *g* for 10 min, and the supernatant (i.e., TritonX-100 soluble fraction) was sampled by adding 2× sample buffer, followed by boiling at 98°C for 3 min to be the Triton-X-soluble fraction. The remaining pellet (i.e., TritonX-100 insoluble fraction) was washed with pellet wash buffer (20 mM Tris-HCl pH7.5, 150 mM NaCl and 10 mM EDTA) using a Vortex-Genie 2 mixer (Scientific Industries). After centrifugation at 4 °C and ∼17,500 × *g* for 10 min, the supernatant was removed, and the pellet was eluted with pellet elusion buffer (20 mM Tris-HCl pH7.5, 150 mM NaCl, 10 mM EDTA, 4% SDS and 20 mM DTT) using a Vortex mixer. The eluted sample was incubated at 65°C for 10 min, followed by sonication for 10 sec twice. After centrifugation at 4 °C and ∼17,500 × *g* for 5 min, the supernatant was sampled by adding 2× sample buffer as the Triton-X-insoluble and SDS-soluble fraction.

The samples were resolved by SDS-PAGE and electroblotted onto a Immobilon-P membrane (Millipore, Cat. #IPVH00010). The membranes were blocked with 3% or 5% skim milk (Megmilk Snow Brand) in TBS-T and probed with the appropriate primary antibodies diluted by the antibody-dilution buffer (each dilution rate is summarized in Supplementary Information). After replacing and probing the appropriate secondary antibodies diluted with skim milk in TBS-T, antibody-antigen complexes were detected on X-ray films (FUJIFILM, Cat. #47410-22167 or #47410-26615) using an enhanced chemiluminescence system (GE Healthcare). The films were converted to digital images using a conventional scanner without any adjustment.

For presentation, representative images were acquired by linearly adjusting the brightness and contrast using Fiji/ImageJ software. When digitally cropping superfluous lanes from blot images, the cropping procedure was executed after the adjustment of brightness and contrast, and cropped position was clearly indicated. Quantification was performed against the raw digital images with densitometry using Fiji/ImageJ software. Kinase activity was defined as the band intensity ratio of phosphorylated protein to total protein. For the kinase activity of endogenous ASK3, the ratio of phosphorylated ASK to ASK3 was calculated.

#### Cell volume measurements

Cell volume changes were measured as the changes in the intracellular water space based on a calcein self-quenching method (Hamann et al., 2002). HeLa cells were seeded in a 96-well plate and incubated for 48 h. Prior to measurements, 40 µM calcein-AM (eBioscence, Cat. #65-0853) was loaded via medium change, and the plate was incubated in 5% CO_2_ at 37°C for 2 h, followed by washing with isoosmotic buffer and replacing with isoosmotic culture medium at 100 µL per well. Before applying hyperosmotic stress, fluorescence (Ex: 485 ± 9 nm, Em: 525 ± 20 nm) was measured in 5% CO_2_ at 37°C for 5min at 1-min intervals using an Infinite 200 Pro. Then, the cells were exposed to hyperosmotic stress by adding 100 µL of 2× hyperosmotic medium per well, and subsequent measurements were performed in 5% CO_2_ at 37°C for 30 min at 1-min intervals.

In the experiments for the removal of hyperosmotic stress, calcein-containing medium was replaced with isoosmotic culture medium at 50 µL per well after washing with isoosmotic buffer. Before applying hyperosmotic stress, fluorescence was measured in 5% CO_2_ at 37°C for 5min at 1-min intervals, and then, the cells were exposed to hyperosmotic stress (600 mOsm) by adding 50 µL of 2× hyperosmotic medium per well, and fluorescence was measured for 20 min. Subsequently, the cells were reverted to isoosmotic conditions by adding 100 µL of ultrapure water per well and observed for another 30 min.

For the data analysis, each fluorescent value was applied background subtraction by the fluorescent value of non-calcein-loaded well, followed by the calculation to the relative fluorescence change to the mean of fluorescent values before hyperosmotic stress, *F*(*t*)/*F*_before_. For the drift correction of volume-independent quenching of calcein and the cellular response against osmotic stress-operation, *F*(*t*)/*F*_before_ was normalized by *F*(*t*)/*F*_before_ of 300 mOsm at each time point.

#### Computational simulation

##### Description of the General Cluster Model

To investigate the intracellular behaviors of biomolecular condensates in silico, we developed a cluster formation model of aggregation-prone proteins in two-dimensional grid space (Dine et al., 2018; Watanabe et al., 2021), referred as “General Cluster Model (GCM)” (Figure 4A). This model contains two types of objects in the square grid space: a protein unit representing an aggregation-prone protein and an obstacle unit representing macromolecular collection which occupies grid space without interacting with other molecules (i.e., as the effect of macromolecular crowding). The consecutive protein units are regarded as a cluster, corresponding to biomolecular condensates in cells. In this model, osmotic cell volume perturbations are achieved by changing the grid space (Watanabe et al., 2021). Each object unit can move to an adjacent grid position according to the following rate constants for the movements of target object (*k*_mov_) (Figure 4A):

- If a target protein unit has no neighboring protein units before the movement, the movements is regarded as simple diffusion with the rate constant *k*_mov1_.
- If a target protein unit has neighboring protein units, unbinding of a target protein unit from other neighboring protein units is regarded as a chemical reaction accompanying breakages of binding relationship. Hence, the rate constant is described as *k*_mov2_ = *A* × exp (− Δ*E* × *n*_lost_ / *θ*) according to the simple Arrhenius equation, where *A* is frequency factor, Δ*E* is an activation energy in one unbinding reaction, *n*_lost_ is the number of neighboring protein units whose binding relationship with the target protein unit will be lost by the movement, and *θ* is a temperature-like constant. This rate constant determines the aggregation-prone property of protein unit.
- If the movement destination of a target protein unit has been already occupied by another protein unit, the movement is regarded as exchange of these protein units, following the rate constant *k*_mov3_. The rate constant is defined from the total timescale (1 / *k*) of the movements of each protein unit (1 / *k*_mov3_ = 1 / *k*_mov2_ + 1 / *k*_mov2_; that is, *k*_mov3_ = *k*_mov2_ / 2).
- The movement of a target obstacle unit is regarded as simple diffusion (without any interaction or exchange processes with neighboring protein and obstacle units), following the rate constant *k*_mov4_.

As exceptions, *k*_mov_ is always 0 in the following situations:

- The movement destination of a target object is a position out of the grid space.
- The movement destination of a target protein unit has been already occupied by an obstacle unit, or vice versa for the case of a target obstacle unit. In other words, the exchange of these units are not allowed because the obstacle units are defined as the effect of macromolecular crowding.

In this simple model, the PPI strength corresponds to the activation energy for breaking one interaction, which can occur in the unbinding and exchange movements of a protein unit; therefore, one can change only Δ*E* to investigate the effects of PPI on the clusters.

##### Description of the ASK3 Dephosphorylation Model

To investigate the relationship between ASK3 condensates and ASK3 dephosphorylation event in silico, we further developed the GCM into a model specific for ASK3 inactivation, referred as “ASK3 Dephosphorylation Model (ADM)” (Figure 5A). This model holds all the definitions of the GCM, with three additional definitions. (1) This model contains two species of protein units, ASK3 and PP6, both of which have aggregation-prone property, corresponding to the protein unit in the GCM. (2) Two states are assigned to the ASK3 unit, phosphorylated (p-ASK3) and dephosphorylated (d-ASK3) states. (3) Reaction events of ASK3 autophosphorylation and dephosphorylation are defined: (3-i) a p-ASK3 unit is dephosphorylated at a certain rate (*k*_rxn1_ or *k*_rxn2_), when the p-ASK3 unit is in the destination of a target PP6 unit upon movements, or vice versa for the inverse target–destination relationship; (3-ii) a d-ASK3 unit is phosphorylated at a certain rate (*k*_rxn3_), when the d-ASK3 unit is in the destination of a target p-ASK3, or vice versa for the inverse target–destination relationship. The interaction between ASK3 unit and PP6 unit is ignored for simplicity. As well as the GCM, each object unit can move to an adjacent grid position according to the following rate movement constants (*k*_mov_) (Figure 5A):

- Free diffusion of a target ASK3 unit or PP6 unit is obeyed at the rate constant *k*_mov1_ or *k*_mov4_, respectively, which is the counterpart of the rate constant *k*_mov1_ in the GCM.
- The rate constant for unbinding of a target ASK3 unit from other neighboring ASK3 units is described as *k*_mov2_ = *A* × exp (− Δ*E*_ASK3–ASK3_ × *n*_ASK3lost_ / *θ*), as well as the rate constant *k*_mov2_ in the GCM, where *A* is frequency factor, Δ*E*_ASK3– ASK3_ is an activation energy for one unbinding reaction between ASK3 units, *n*_ASK3lost_ is the number of neighboring ASK3 units whose binding relationship with the target ASK3 unit will be lost by the movement, and *θ* is a temperature-like constant. Likewise, the rate constant for unbinding of PP6 unit from other neighboring PP6 units is described as *k*_mov5_ = *B* × exp (− Δ*E*_PP6–PP6_ × *n*_PP6lost_ / *θ*).
- The rate constant for exchange of ASK3 units is defined as 1 / *k*_mov3_ = 1 / *k*_mov2_ + 1 / *k*_mov2_; that is, *k*_mov3_ = *k*_mov2_ / 2, as well as the rate constant *k*_mov3_ in the GCM. Likewise, the rate constant for the exchange of PP6 units is defined as 1 / *k*_mov6_ = 1 / *k*_mov5_ + 1 / *k*_mov5_; that is, *k*_mov6_ = *k*_mov5_ / 2.
- The rate constant for exchange between ASK3 monomer unit (without neighboring ASK3 units) and PP6 monomer unit (without neighboring PP6 units) is defined from the total timescale (1 / *k*) of the free diffusions of each unit (1 / *k*_mov7_ = 1 / *k*_mov1_ + 1 / *k*_mov4_; that is, *k*_mov7_ = *k*_mov1_ × *k*_mov4_ / (*k*_mov1_ + *k*_mov4_)).
- The movement of a target obstacle unit is regarded as simple diffusion with the rate constant *k*_mov8_, which is the counterpart of the rate constant *k*_mov4_ in the GCM.

As exceptions, *k*_mov_ is set to be 0 in the following situations:

- The movement destination of a target object is out of grid space, consistent with the GCM.
- An obstacle unit exists in the movement destination of a target protein unit, or vice versa, consistent with the GCM.
- When either target or destination ASK3 unit has neighboring ASK3 units or either target or destination PP6 unit has neighboring PP6 units, exchange between ASK3 unit and PP6 unit was not allowed based on the observation that ASK3 condensates and ANKRD52 condensates exists in each own phase (regardless TRPM4 inhibition) at the early time points after hyperosmotic stress when the dephosphorylation of ASK3 has been accomplished (Figure 2D and S3).

The reaction rate constant *k*_rxn_ is designated as follows (Figure 5A):

- To analyze the significance of the condensate formation in ASK3 inactivation, we prepare rate constants for ASK3 dephosphorylation in or out of ASK3 cluster separately. When a PP6 unit is in the destination of a target p-ASK3 unit outside of ASK3 cluster, or vice versa for the inverse target–destination relationship, the dephosphorylation reaction obeys the rate constant *k*_rxn1_. When a PP6 unit is in the destination of a target p-ASK3 in ASK3 cluster, or vice versa, the dephosphorylation obeys the rate constant *k*_rxn2_.
- When a d-ASK3 unit is in the movement destination of a target p-ASK3 unit, or vice versa, the autophosphorylation reaction obeys the rate constant *k*_rxn3_.

In this model, the ratio of p-ASK3 units to total ASK3 (p-ASK3 + d-ASK3) units corresponds to the ASK3 activity of a cell. To investigate the effects of PPI on the ASK3 activity, one can change Δ*E*_ASK3–ASK3_ in *k*_mov2_ and *k*_mov3_ independently.

##### Simulation of the General Cluster Model

To simulate the GCM in silico, we computed random trajectories of protein units and obstacle units in two dimensional square grid space using rejection kinetic Monte Carlo method in Pyhton3 language, as previously described (Watanabe et al., 2021). Simulations of our computational models were performed on the supercomputer SHIROKANE (Human Genome Center, Institute of Medical Science, University of Tokyo), which enabled parallel executions while covering sufficiently large parameter space and trials (Niida et al., 2019). Simulation algorithm was executed as follows.

Step 1: A target unit with a chance to move was randomly picked from the union of protein units and obstacle units, followed by the random selection of a potential destination of the target unit from 8 adjacent candidates.

Step 2: Rate constant of the movement (*k*_mov_) was systemically calculated from the designation of the target unit and the destination (see Description of the model).

Step 3: A random number *r* was acquired from the interval [0,1).

Step 4: If *k*_mov_ > *r*, the movement was accepted: the target unit was renewed at the position of destination. Otherwise, if *k*_mov_ ≤ *r*, the movement was rejected: the target unit stayed at the original position.

A series of procedures from step 1 to step 4 was regarded as one iteration, and we executed 1 × 10^7^ iterations.

For the simulation of GCM (Figure 4), we prepared 500 protein units and 1,500 obstacle units randomly located in the grid space with 65 × 65 squares as the initial condition. Six consecutive protein units were defined as the minimum size of the cluster. We set *k*_mov1_ = 1; therefore, free diffusion of ASK3 units was always accepted. In contrast, we set *k*_mov4_ = 0.01; therefore, the diffusion of obstacle units was slower than diffusion of ASK3 units, because the movement of an obstacle unit virtually represents the average movements of the constituent macromolecules, whose movement vectors mostly cancel each other out in a real cell. For the penalty defining rate constant *k*_mov2_, we set *A* = 1 to adjust *k*_mov2_ = 1 under the condition where *n*_lost_ = 0, and fixed *θ* = 1 for simplicity. We practically ignored *k*_mov3_ in our Phython3 scripts because the exchange event between protein units does not distinguish their positions regardless of whether it actually happens or not. To investigate the effect of PPI on the morphology of cluster, Δ*E* was changed ranging from 1 to 4.

##### Simulation of the ASK3 Dephosphorylation Model

As the GCM simulation in silico, we simulated the ADM using rejection kinetic Monte Carlo method in Pyhton3 language with SHIROKANE (Human Genome Center, Institute of Medical Science, University of Tokyo). Simulation algorithm was executed as follows.

First, only the movements of object units were executed to develop cluster formation as same as the simulation of the GCM, except that a target unit was randomly picked from the union of ASK3 units, PP6 units and obstacle units. After this initial cluster formation, subsequent main iterations included the following reaction steps 5–7 for dephosphorylation or autophosphorylation event, in addition to the movement procedures from step 1 to step 4 (see Simulation of the General Cluster Model).

Step 5: Rate constant of the reaction (*k*_rxn_) was designated according to the target (ASK3 or PP6) unit and the destination selected at step 1.

Step 6: Another random number *r*’ was acquired from the interval [0,1).

Step 7: If *k*_rxn_ > *r*’, the reaction was accepted: the phosphorylation state of the ASK3 unit that involves in the reaction (i.e., target unit or the unit located in destination) was changed to the other state. Otherwise, if *k*_rxn_ ≤ *r*’, the reaction was rejected.

That is, the movements and the reaction were independently calculated in one iteration. We executed 5 × 10^6^ iterations for the initial cluster formation and 5 × 10^6^ or 2 × 10^6^ iterations in Figure 5 or Figure S4A, respectively, for main iterations.

For the simulation of the ADM (Figure 5), we prepared 500 p-ASK3 units, 500 PP6 units and 1,500 obstacle units randomly located in the grid space with 70 × 70 squares as the initial condition. Six consecutive ASK3 or PP6 units were defined as the minimum size of the ASK3 or PP6 cluster, respectively. We set *k*_mov1_ = *k*_mov4_ = 1; therefore, free diffusion of ASK3 units or PP6 units was always accepted. In contrast, we set *k*_mov8_ = 0.01; therefore, the diffusion of obstacle units was slower than diffusion of ASK3 units or PP6 units. For the penalty-defining rate constant *k*_mov2_ and *k*_mov5_, we set *A* = 1 to adjust *k*_mov2_ = 1 under the condition where *n*_ASK3lost_ = 0, and set *B* = 1 to adjust *k*_mov4_ = 1 under the condition where *n*_PP6lost_ = 0. *θ* was fixed at 1 for simplicity. We practically ignored *k*_mov6_ in our Phython3 scripts, because PP6 units are defined to have no status information and thus the exchange event between PP6 units does not distinguish their positions regardless of whether it actually happens or not. PPI defining parameter Δ*E*_ASK3–ASK3_ in *k*_mov2_ and *k*_mov3_ was independently changed ranging from 1 to 4. Δ*E*_PP6–PP6_ was fixed at 2 to be stronger than the default Δ*E*_ASK3–ASK3_ (= 1), because the FRAP of ANKRD52 condensates was slower than that of ASK3 condensates and was not affected by intracellular Na^+^ concentration (Figure S4B and S4C). For the rate constants designated to dephosphorylation reaction events, we set the same values between them (*k*_rxn1_ = *k*_rxn2_ = 0.1) in Figure S4A and different values (*k*_rxn1_ = 0, *k*_rxn2_ = 0.1) in Figure 5 to investigate the importance of ASK3 cluster in ASK3 dephosphorylation. For autophosphorylation, we set to be *k*_rxn3_ = 0.001. Note that it is not unusual to assume that dephosphorylation is faster than autophosphorylation, because phosphorylation accompanies three molecules, kinase, substrate and ATP, whereas dephosphorylation does two, phosphatase and substrate. The parameter values of these reaction rate constants (*k*_rxn_) were determined from pilot simulation.

##### Summary of simulation results

For figure presentation, our Python3 scripts saved the coordinates and species of each molecule, which was rendered using RStudio software. For time-lapse video, a series of images with added captions were converted to a movie file using Fiji/ImageJ software. For quantification, our Python3 script calculated the count/mean size of cluster and ASK3 activity (i.e., the ratio of p-ASK3 units to total ASK3 (p-ASK3 + d-ASK3) units). The exported data table was summarized in RStudio software.

#### Quantification and statistical analysis

The data are summarized as the mean ± SEM except for boxplots (center line = median; box limits = [*Q*_1_, *Q*_3_]; whiskers = [max(minimum value, *Q*_1_ − 1.5 × IQR), min(maximum value, *Q*_3_ + 1.5 × IQR)], where *Q*_1_, *Q*_3_ and IQR are the first quartile, the third quartile and the interquartile range, respectively). No statistical method was utilized to predetermine the sample size because all experiments in this study were performed with defined laboratory reagents and cell lines. Based on several pilot experiments to determine the experimental conditions, each sample size was chosen as large as possible to represent experimental variation while still practically feasible in terms of data collection. All the statistical tests were performed using RStudio software, and *P* < 0.05 was considered statistically significant. Each statistical method was described in figure legend.

## Acknowledgements

We thank Y. Hata (Tokyo Medical and Dental University) and J. Maruyama (RIKEN Center for Integrative Medical Sciences) for kindly providing the TAZ expression plasmid, A. Kakizuka (Kyoto University) for kindly providing the polyQ expression plasmids, and S. Uchida (Tokyo Medical and Dental University) for kindly providing the WNK1 expression plasmid. This work was supported by the Japan Agency for Medical Research and Development (AMED) under the Project for Elucidating and Controlling Mechanisms of Aging and Longevity (grant number JP21gm5010001 to H.I.), by the Japan Society for the Promotion of Science (JSPS) under the Grants-in-Aid for Scientific Research (KAKENHI; grant numbers JP17K15086 to K.W. JP18H02569 to I.N. and JP18H03995 and JP21H04760 to H.I.) and by the Japan Science and Technology Agency (JST) Moonshot R&D— MILLENNIA Program (grant number JPMJMS2022-18 to H.I.).

## Author contributions

K.W., I.N. and H.I. conceptualized and supervised this project. K.M. designed and performed all experiments, computational simulations, statistical analyses, data visualizations and figure illustrations. K.W. established assay systems for condensation in cell and in vitro. K.M., K.W., I.N. and H.I. wrote and revised the manuscript.

