## Supplementary Materials for "Sodium ion regulates liquidity of biomolecular condensates in hyperosmotic stress response"

**This PDF file includes:**

Figure S1 to S6
Captions for Supplementary Videos
Key Resource Table

**Other Supplementary Materials for this manuscript include the following:**

Video S1 to S3

### Supplementary Figures

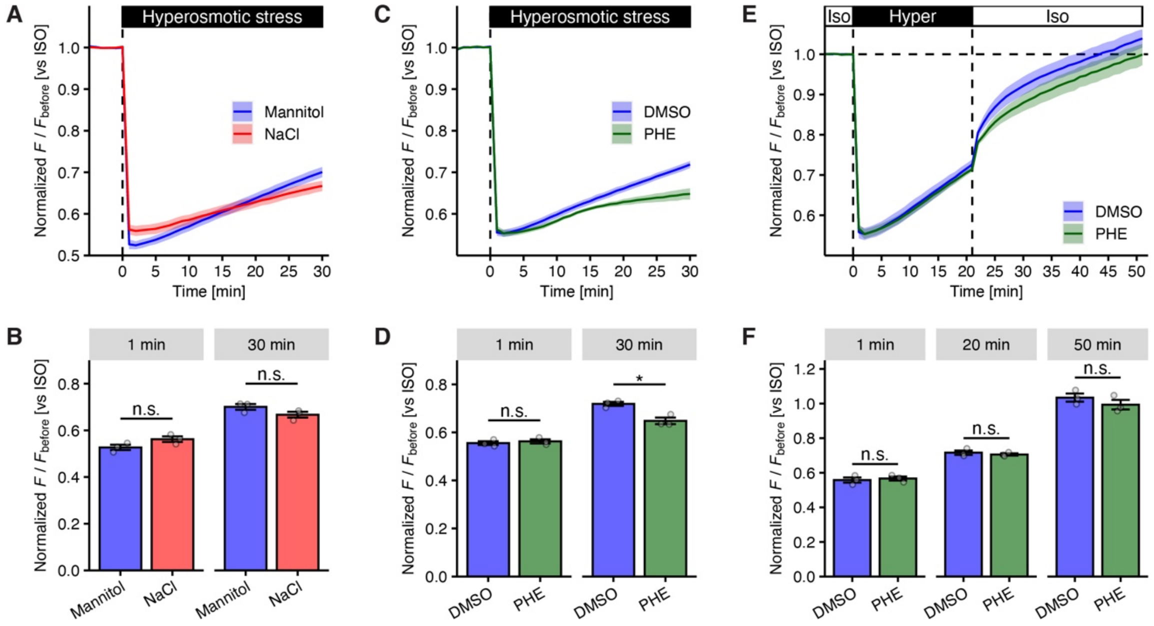


**Figure S1 Cell volume recovery from the cell shrinkage induced by hyperosmotic stress.**
(A, B) Cell volume recovery under hyperosmotic stress. Changes in cell volume over time (A) and the cell volume at the indicated time points (B) in HeLa cells are presented. Hyperosmotic stress: mannitol- or NaCl-supplemented medium (600 mOsm).
(C, D) Cell volume recovery under TRPM4 inhibition. Changes in cell volume over time (C) and the cell volume at the indicated time points (D) in HeLa cells are presented. Hyperosmotic stress: mannitol-supplemented medium (600 mOsm).
(E, F) Cell volume changes following the removal of hyperosmotic stress. Changes in cell volume over time (E) and the cell volume at the indicated time points (F) in HeLa cells are presented.
DMSO: solvent for 9-phenanthrol. PHE: 20 μM (C, D) or 15 μM (E, F) 9-phenanthrol (TRPM4 inhibitor). These reagents were treated at the same time as the hyperosmotic stress. Cell volume was measured with a calcein-quenching assay in HeLa cells. The Y-axis represents the normalized value of calcein fluorescence ($F$), which correlates with cell volume. Data: mean ± SEM; $n$ = 3 independent experiments; n.s. (not significant), $^{*}P < 0.05$, according to two-sided unpaired Student's $t$-tests.

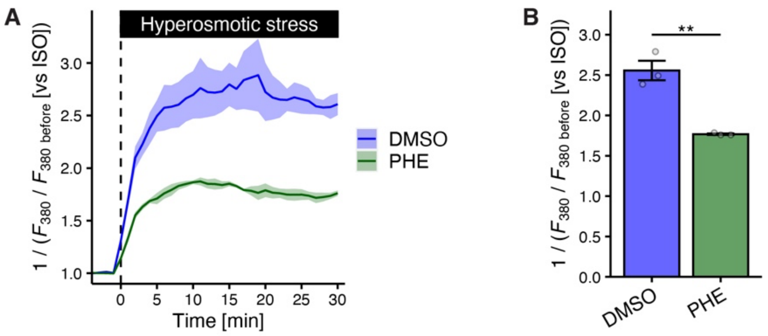


**Figure S2 TRPM4 serves $Na^+$ influx under hyperosmotic stress**

$[Na^+]_i$ under hyperosmotic stress with TRPM4 inhibition. Changes in $[Na^+]_i$ following hyperosmotic stress over time (A) and $[Na^+]_i$ at 30 min (B) in HeLa cells are presented. $F_{380}$ represents fluorescence from SBFI excited at 380 nm, and the Y-axis correlates with $[Na^+]_i$. Hyperosmotic stress: mannitol-supplemented medium (600 mOsm). DMSO: solvent for 9-phenanthrol. PHE: 25 μM 9-phenanthrol (TRPM4 inhibitor). Data were smoothed by the moving average of 3 time points. Data: mean ± SEM; $n$ = 3 independent experiments. $^{**}P < 0.01$ according to two-sided unpaired Student's $t$-tests.

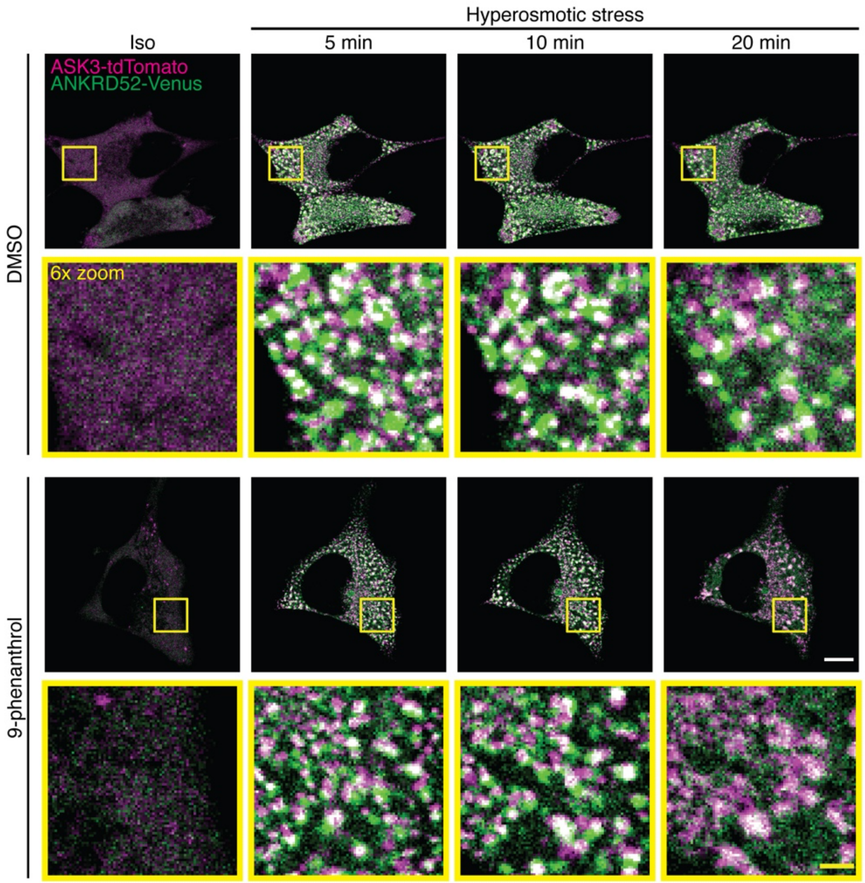


**Figure S3 ASK3 condensates lost the phase boundary with PP6 condensates under TRPM4 inhibition in the late phase of hyperosmotic stress.**

Representative images of the relationship between ASK3 condensates and ANKRD52 condensates in HEK293A cells under mannitol-supplemented hyperosmotic stress (550 mOsm). Magenta: ASK3-tdTomato, green: ANKRD52-Venus. DMSO: solvent for 9-phenanthrol. PHE: 20 µM 9-phenanthrol (TRPM4 inhibitor). Scale bar: 10 µm (white) or 2 µm (yellow). Note that the signal intensity cannot be compared between samples.

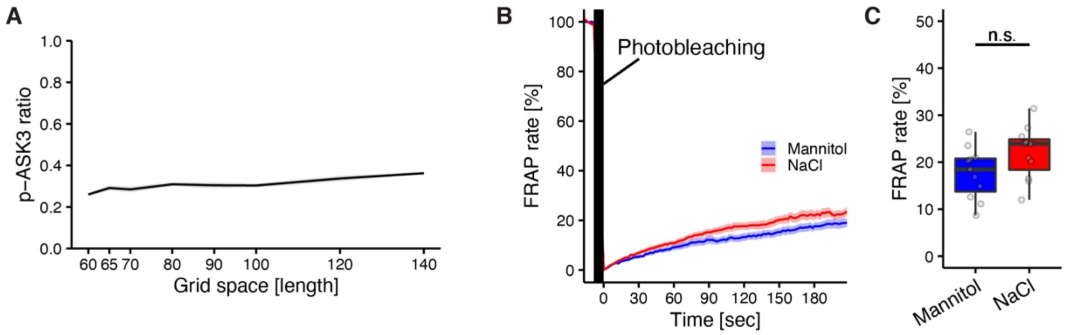


**Figure S4 Determination of parameter values in the simulation of ASK3 inactivation.**
(A) A computational simulation of the relationship between the grid space and the ratio of p-ASK3 to total ASK3 under the condition that ASK3 is dephosphorylated inside and outside ASK3 cluster at the same reaction rate. The results after 5 × $10^6$ iteration steps for cluster formation and another 2 × $10^6$ iteration steps for dephosphorylation are presented. Data: mean ± SEM, $n$ = 6 simulations. (B, C) FRAP assay of ANKRD52 condensates. Changes in the FRAP of ANKRD52 condensates over time (B) and the FRAP at 180 sec (C) in ANKRD52-tdTomato-transfected HEK293A cells are presented. Hyperosmotic stress: mannitol- or NaCl-supplemented medium (600 mOsm). Data: mean ± SEM (B) or centerline = median; box limits = [$Q_1$, $Q_3$]; whiskers = [max(minimum value, $Q_1$ − 1.5 × IQR), min(minimum value, $Q_3$ + 1.5 × IQR)], where $Q_1$, $Q_3$ and IQR are the first quartile, the third quartile and the interquartile range, respectively (C). $n$ = 11 cells pooled from 3 independent experiments. n.s. (not significant), according to a two-sided Welch's $t$-test.

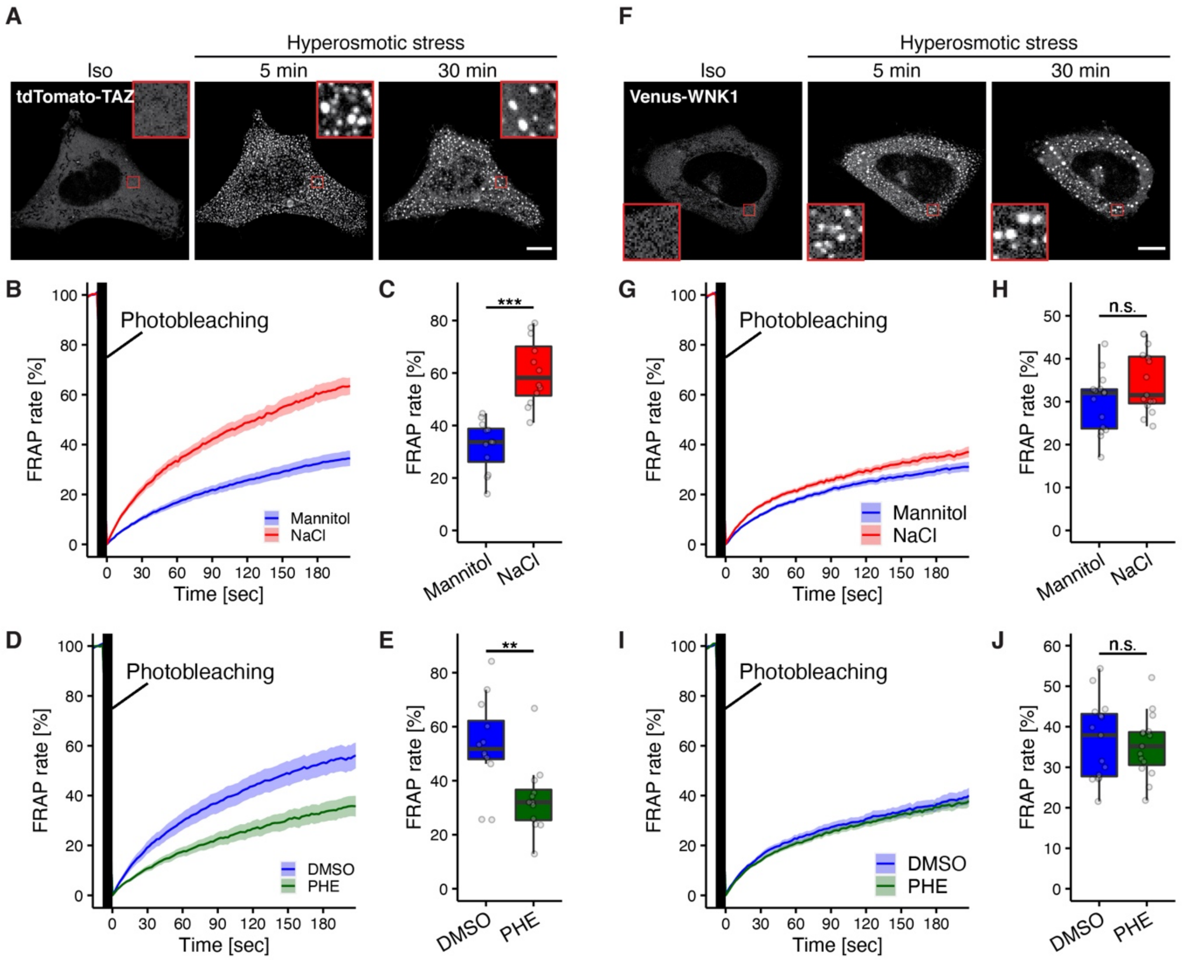


**Figure S5 The effects of intracellular $Na^+$ on the liquidity of TAZ condensates or WNK1 condensates.**
(A) Representative images of TAZ condensates under hyperosmotic stress. tdTomato-TAZ was transfected into HEK293A cells. Hyperosmotic stress: mannitol-supplemented medium (550 mOsm). (B–E) FRAP assay of TAZ condensates. Changes in the FRAP of TAZ condensates over time (B, D) and the FRAP at 180 sec (C, E) in tdTomato-TAZ-transfected HEK293A cells are presented. Hyperosmotic stress: mannitol- or NaCl-supplemented medium (700 mOsm) (B, C), or mannitol-supplemented medium (550 mOsm) under TRPM4 inhibition (D, E). PHE: 20 µM of 9-phenanthrol treated at the same time as osmotic stress.
(F) Representative images of WNK1 condensates under hyperosmotic stress. Venus-WNK1 was transfected into HEK293A cells. Hyperosmotic stress: mannitol-supplemented medium (550 mOsm). (G–J) FRAP assay of WNK1 condensates. Changes in the FRAP of WNK1 condensates over time (G, I) and the FRAP at 180 sec (H, J) in tdTomato-WNK1-transfected HEK293A cells are presented. Hyperosmotic stress: mannitol- or NaCl-supplemented medium (600 mOsm) (B, C), or mannitol-supplemented medium (550 mOsm) under TRPM4 inhibition (D, E). PHE: 25 µM of 9-phenanthrol treated at the same time as osmotic stress.
(A, F) Scale bar: 10 µm. Red square: 5 × zoom. (B, D, G, I) Data: mean ± SEM. (C, E, H, J) Data: centerline = median; box limits = [$Q_1$, $Q_3$]; whiskers = [max(minimum value, $Q_1 - 1.5 \times$ IQR), min(minimum value, $Q_3 + 1.5 \times$ IQR)], where $Q_1$, $Q_3$ and IQR are the first quartile, the third quartile and the interquartile range, respectively. $n = 12$ (B–E) or 15 (G–J) cells pooled from 3 independent experiments. $^{**}P < 0.01$, $^{***}P < 0.001$, n.s. (not significant), according to two-sided Welch's $t$-tests.

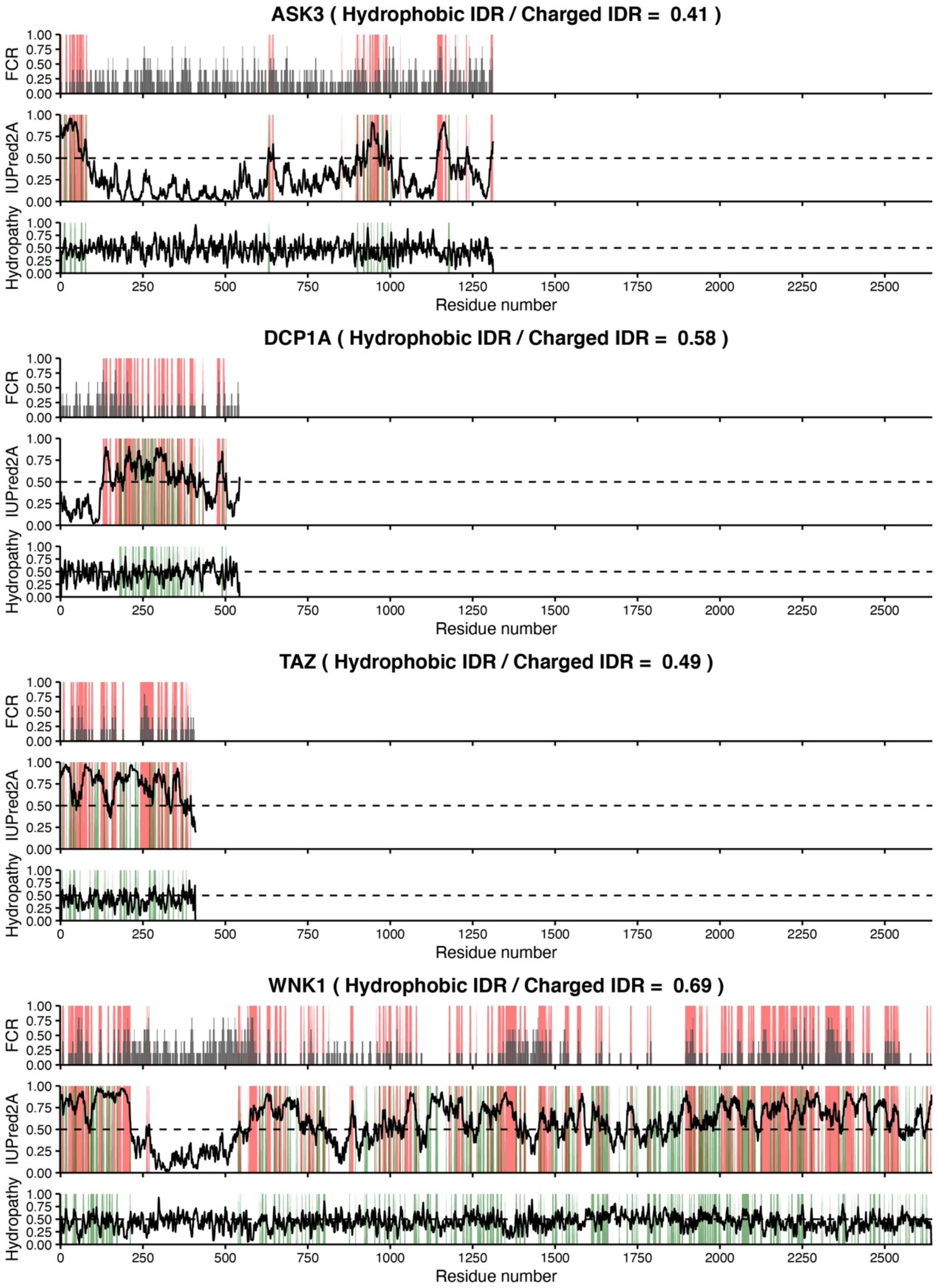


**Figure S6 Analysis of amino acid sequence composition.**

Characterization of amino acid sequences of ASK3, DCP1A, TAZ and WNK1. Red: charged IDRs defined as IDRs whose fraction of charged residues (FCR) was more than 0; green: hydrophobic IDRs defined as IDRs whose hydropathy was 0.5 or more. The ratio of "hydrophobic IDRs/charged IDRs" represents the ratio of the number of residues colored green to those colored red. The FCR and hydropathy of the amino acid sequence were calculated using localCIDER in Python3 (Holehouse et al., 2017; Kyte and Doolittle, 1982). The IDRs were predicted using the IUPred2A tool (Mészáros et al., 2018). A sliding window contained 5 residues.

**Supplementary Videos**

**Video S1 Dynamics and fusion of ASK3 condensates under mannitol- or NaCl-induced hyperosmotic stress**

After 5 min, the cells were exposed to mannitol- or NaCl- induced hyperosmotic stress (600 mOsm). Scale bar: 10 μm.

**Video S2 A computational simulation for the relationship between protein–protein interaction and cluster fluidity.**

A representative video of the GCM simulation (See the STAR Methods). Each frame was representative image at every 20,000 iteration steps. Red square: protein unit, black square: obstacle unit.

**Video S3 A computational simulation for ASK3 dephosphorylation.**

A representative video of the ADM simulation (See the STAR Methods). ASK3 dephosphorylation steps after cluster formation. Each frame was representative image at every 10,000 iteration steps. Red square: p-ASK unit, orange square: d-ASK3 unit, blue square: PP6 unit, black square: obstacle unit. $\Delta E_{\mathrm{ASK3-ASK3}} = 1$ in $k_2$ and $k_3$.

**Key resources table**

| REAGENT or RESOURCE | SOURCE | IDENTIFIER |
|---|---|---|
| Antibodies | | |
| Anti-rabbit IgG, HRP-linked (IB: 1:1,000–1:20,000) | Cell Signaling Technology | Cat. #7074 |
| Anti-rat IgG, HRP-linked (IB: 1:5,000–1:10,000) | Cell Signaling Technology | Cat. #7077 |
| Anti-mouse IgG, HRP-linked (IB: 1:1,000–1:20,000) | Cell Signaling Technology | Cat. #7076 |
| Rabbit polyclonal anti-phospho-ASK (p-Ask; Thr808 in human ASK3 and Thr838 in human ASK1, IB: 1:10,000 or 1:1,000) | Naguro et al., 2012; Tobiume et al., 2002 | N/A |
| Mouse monoclonal anti-DYKDDDDK tag (FLAG; clone 1E6, IB: 1:10,000) | Wako Pure Chemical Industries | Cat. #012-22384 |
| Mouse monoclonal anti-Actin (Actin; clone AC-40, IB: 1:2,000) | Sigma-Aldrich | Cat. #A3853 |
| Rabbit monoclonal anti-ASK1 (ASK1; clone EP553Y, IB: 1:10,000) | Abcam | Cat. #ab45178 |
| Rat monoclonal anti-ASK3 (ASK3; IB: 1:10,000) | Naguro et al., 2012 | N/A |
| Mouse monoclonal anti-TRPM4 (TRPM4; clone OTI10H5, IB: 1:1,000) | Abcam | Cat. #ab123936 |
| Mouse monoclonal anti-GFP (GFP; clone 1E4, IB: 1:10,000) | Medical & Biological Laboratories | Cat. #M048-3 |
| Rat monoclonal anti-α-Tubulin (α-Tubulin; clone YL1/2, IB: 1:10,000) | Santa Cruz Biotechnology | Cat. #sc-53029 |
| Rabbit polyclonal anti-H3 (H3, IB: 1:50,000) | Abcam | Cat. #ab1791 |
| ANTI-FLAG M2 Affinity Gel (clone M2, IP) | Sigma-Aldrich | Cat. #A2220 |
| Chemicals, peptides, and recombinant proteins | | |
| Polyethyleneimine "MAX" | Polyscience | Cat. #24765 |
| Lipofectamine RNAiMAX | Invitrogen | Cat. #133778-150 |
| Lipofectamine 2000 Transfection Reagent | Invitrogen | Cat. #11668019 |
| Zeocin | Invitrogen | Cat. #R25001 |
| Blasticidin S HCl | Invitrogen | Cat. #A1113903 |
| Tetracycline | Sigma-Aldrich | Cat. #T7660 |
| 3x FLAG peptide lyophilized powder | Sigma-Aldrich | Cat. #F4799 |
| *N*-(3-trifluoromethylphenyl)anthranilic acid (flufenamate) | Tokyo Chemical Industry | Cat. #T2354 |
| 9-phenanthrol | Sigma-Aldrich | Cat. #211281 |
| Calcein-AM | eBioscience | Cat. #65-0853 |
| SBFI-AM | Invitrogen | Cat. #S1263 |
| Pluronic F-127 | Invitrogen | Cat. #P3000MP |
| 1,6-hexanediol | Wako Pure Chemical Industries | Cat. #081-00435 |
| Polyethylene glycol 4000 | Kanto Kagaku | Cat. #32828-02 |
| N-Methyl-D(−)-glucamine (NMDG) | Wako Pure Chemical Industries | Cat. #132-08012 |
| Experimental models: Cell lines | | |
| Human: HEK293A cells | Invitrogen | N/A |
| Human: Tetracycline-inducible Flag-ASK3-stably-expressing HEK293A cells | Watanabe et al., 2018 | N/A |
| Human: Tetracycline-inducible Venus-ASK3-stably-expressing HEK293A cells | Watanabe et al., 2021 | N/A |
| Human: HeLa cells | ATCC | N/A |
| Oligonucleotides | | |

| | | |
|---|---|---|
| Control siRNA #1 (ON-TARGET*plus* Non-targeting siRNA #1) | Horizon | Cat. #D-001810-01 |
| Control siRNA #2 (ON-TARGET*plus* Non-targeting siRNA #2) | Horizon | Cat. #D-001810-02 |
| TRPM4 siRNA #1 (ON-TARGET*plus* Human TRPM4 siRNA #1) | Horizon | Cat. #J-006515-09 |
| TRPM4 siRNA #2 (ON-TARGET*plus* Human TRPM4 siRNA #2) | Horizon | Cat. #J-006515-10 |
| TRPM4 siRNA #3 (ON-TARGET*plus* Human TRPM4 siRNA #3) | Horizon | Cat. #J-006515-11 |
| TRPM4 siRNA #4 (ON-TARGET*plus* Human TRPM4 siRNA #4) | Horizon | Cat. #J-006515-12 |
| Recombinant DNA | | |
| pcDNA4/TO EGFP-FLAG-ASK3 (CDS of NM_001001671.3 with c.147C>T, c.574G>A) | Watanabe et al., 2021 | N/A |
| pcDNA3 ASK3-tdTomato (CDS of NM_001001671.3 with c.147C>T, c.574G>A) | Watanabe et al., 2021 | N/A |
| pcDNA3/GW ANKRD52-Venus (CDS of NM_173595.3) | Watanabe et al., 2021 | N/A |
| pcDNA3/GW Venus-DCP1A (CDS of NM_018403.7) | Watanabe et al., 2021 | N/A |
| pcDNA3/GW tdTomato-DCP1A (CDS of NM_018403.7) | This paper | N/A |
| pcDNA3/GW tdTomato-TAZ (CDS of NM_001168278.3) | This paper (cDNA from Dr. Hata) | N/A |
| pcDNA3/GW Venus-WNK1 (CDS of NM_018979.4 with c.258T>C, c.1053A>G, c.1287A>G, c.2001T>A, c.2002C>A, c.2328G>A, c.3147G>A, c.3166A>C, c.3690A>G, c.4044C>T, c.4517G>C, c.6828C>T, c.6906T>C, c.6828C>T, c.6909T>C) | This paper (cDNA from Dr. Uchida) | N/A |
| pcDNA3/GW tdTomato-WNK1 (CDS of NM_018979.4 with c.258T>C, c.1053A>G, c.1287A>G, c.2001T>A, c.2002C>A, c.2328G>A, c.3147G>A, c.3166A>C, c.3690A>G, c.4044C>T, c.4517G>C, c.6828C>T, c.6906T>C, c.6828C>T, c.6909T>C) | This paper (cDNA from Dr. Uchida) | N/A |
| pcDNA4/TO Venus-polyQ34 (Q3+K+Q31) | This paper (cDNA from Dr. Kakizuka) | N/A |
| pcDNA4/TO Venus-polyQ71 | This paper (cDNA from Dr. Kakizuka) | N/A |
| Software and algorithms | | |
| Language: Python3 (ver. 3.8.2, ver.3.6.5) | Python Software Foundation | https://www.python.org/ |
| Python library: Numpy (ver. 1.18.4, ver. 1.15.2) | NumPy Developers | https://numpy.org/ |
| Python library: Pandas (ver. 1.0.3) | PyData Development Team | https://pandas.pydata.org/ |
| Python library: Matplotlib (ver. 3.2.1) | Hunter et al. | https://matplotlib.org/ |
| Python library: localCIDER (ver. 0.1.19) | Holehouse et al. 2015 | http://pappulab.github.io/localCIDER/ |
| Language: R (ver. 4.0.3) | R Foundation | https://www.r-project.org/ |
| RStudio (ver. 1.2.5042) | RStudio | https://www.rstudio.com/ |
| R package: tidyverse (ver. 1.3.0) | H. Wickham (RStudio team) | https://www.tidyverse.org/ |
| R package: ggpubr (ver. 0.4.0) | A. Kassambara | https://rpkgs.datanovia.com/ggpubr/ |
| R package: RColorBrewer (ver. 1.1.2) | E. Neuwirth | https://cran.r-project.org/package=RColorBrewer |
| R package: multcomp (ver. 1.4.15) | T. Hothorn et al. | http://multcomp.R-forge.R-project.org |

| | | |
|---|---|---|
| R package: zoo (ver. 1.8.8) | A. Zeileis | https://cran.r-project.org/package=zoo |
| ExPASy ProtPram tool | Gasteiger et al. | https://web.expasy.org/protparam/ |
| IUPred2A | Mészáros et al., 2018 | https://iupred2a.elte.hu/ |
| Fiji/ImageJ (ver. 2.3.0) | Schindelin et al., 2012 | https://fiji.sc/ |
| Fiji Plugin: TrackMate (ver. 7.6.1) | Tinevez et al., 2017 | https://imagej.net/TrackMate |
| Other | | |
| 35 mmφ glass bottom dish | Matsunami | Cat. #D11130H |
| 15×24 mm cover slip | Matsunami | Cat. #C01824 |
| 96-well black plate | BM Bio | Cat. #215006 |
